# RegionScan: A comprehensive R package for region-level genome-wide association testing with integration and visualization of multiple-variant and single-variant hypothesis testing

**DOI:** 10.1101/2024.03.04.582374

**Authors:** Myriam Brossard, Delnaz Roshandel, Kexin Luo, Fatemeh Yavartanoo, Andrew D. Paterson, Yun J. Yoo, Shelley B. Bull

## Abstract

**Summary:** RegionScan is an R package for comprehensive and scalable genome-wide association testing of region-level multiple-variant and single-variant statistics and visualization of the results. It implements various state-of-the-art region-level tests to improve signal detection under heterogeneous genetic architectures and facilitates comparison of multiple-variant region-level and single-variant test results. It exploits local linkage disequilibrium (LD) structure for genomic partitioning and LD-adaptive region definition. RegionScan is compatible with VCF input file formats for genotyped and imputed variants, and options are available for analysis of multi-allelic variants and unbalanced binary phenotypes. It accommodates parallel region-level processing and analysis to improve computational time and memory efficiency and provides detailed outputs and utility functions to assist results comparison, visualization, and interpretation.

**Availability and implementation:** RegionScan is freely available for download on GitHub (https://github.com/brossardMyriam/RegionScan).

**Supplementary information:** Supplementary data are available at Bioinformatics online.

## 1. Introduction

Compared to genome-wide single-variant testing, region-level multi-variant association analysis can better capture signals under complex genetic architectures^1^. Because fewer tests are conducted, multiple testing is reduced, and the genome-wide testing threshold can be relaxed. However, for comprehensive genomic analysis, region-level testing requires appropriate region definition, e.g. including intergenic, intronic, and exonic variants. It also faces analytical challenges, including high dimensionality and multi-collinearity within regions produced by complex and long-range linkage disequilibrium (LD) structure. Available region-level tests differ according to the underlying assumptions, the construction of the test statistic, and thus are sensitive to different regional genetic architectures^2,3^. We focus on three classes of state-of-the-art region-level tests (**<u>Supplementary Information 1</u>**), including multi-variant linear/logistic regression tests with and without dimension reduction^3–5^, variance component score tests^6,7^, and region-level min*P* tests^8–10^; sensitive to heterogeneous regional architectures. Our goal is to integrate region definition with implementation of region-level and single-variant tests in one scalable R package for comprehensive genome-wide region-level association analysis and improve region discovery under heterogeneous regional architectures.

## 2. Implementation and Key features

We introduce the RegionScan R package for genome-wide discovery analysis and define regions using the gpart^11^ R package for LD-based genomic partitioning, optimized for region-level analysis^12^. Although a major advantage of our approach is comprehensive analysis of the genome, including intergenic regions, RegionScan can also accommodate other user-specified region definitions.

### 2.1 Capability and Scalability

The main function is called *regscan* (**<u>Supplementary Information 2</u>**, for a detailed description and list of options). *regscan* takes four main inputs: *data* which includes genotypes (additively coded); *SNPinfo* input with variant information; *phenocov* with phenotypes (quantitative or binary) as well as covariates (if applicable) and a *REGIONinfo* input with region start/end positions, as produced with gpart^11^. Alternatively, the auxiliary function *recodeVCF* can process large VCF 4.0 files with vcftools^13^ to produce a temporary subset VCF file by region, subsequently processed in R to improve memory efficiency. *regscan* also deals optionally with multi-allelic variants in addition to bi-allelic variants.

To improve scalability, *regscan* can process, recode and analyze each region in parallel. The processing steps for each region include variant filtering and recoding based on minor allele frequency (MAF), and an option to reduce multicollinearity by pruning variants within regions. This is followed by application of region-level tests including regression-based tests (MLC^2^, PC80^3^, LC^4,5,14,15^, generalized Wald tests), variance component score tests (SKAT^16,17^, SKAT-O^7^), and region-level min *P* tests (simpleM^8^, GATES^9^, MinP^10,18^), in addition to single-SNP tests for variants within regions. *regscan* includes an option to reduce finite-sample bias in logistic regression of unbalanced binary traits and/or variants with low minor allele counts, using a Jeffreys-prior penalized likelihood^19,20^. For the MLC^2^ region-level test, variants within each region are clustered in LD bins based on pairwise correlation^21^ for reduced-*df* region-level testing adaptive to the number of LD bins, followed by variant recoding within each bin to maximize variant pairs positive correlation (**<u>Supplementary Information 2</u>**, section 2.1.1); bin-level tests within regions are reported in addition to MLC region-level tests.

### 2.3 Detailed Outputs and Visualization

*regscan* produces six outputs detailing results for all regions analyzed (**<u>Supplementary Information 2</u>**, section 2.1.3): (1) *region-level* output with results for all regions analyzed; (2) *bin-level* output including bin-level test results for all bins within each region; (3) *variant-level* output with variant positions, LD-bin assignments, and corresponding effect sizes and *P*-values from single- and multi-variant regional regression models; within-region variance inflation factor values (VIFs) are included to facilitate identification of multi-collinearity; (4) a *list of variants pruned out* with reasons for exclusions; and optionally, (5) a s*ingle-variant* output including variant-to-LD bin assignments for all the variants (available before pruning) and (6) a *covariate* output with covariate effects and *P*-values extracted from multi-variant regional regression models.

Utility functions are implemented to visualize comparisons between region and/or variant-level test results. For example, *MiamiPlot* produces a genome-wide comparison of -log10 *P*-values for a pair of tests; *LocusPlot* displays results of several region-level tests in a set of contiguous regions; *QQPlot* assesses consistency of the observed distribution of a specified region-level statistic *P*-value with that expected under the null hypothesis and returns corresponding genomic inflation factors. *regscan* also produces optional heatmap plots within each of a set of selected region(s) to visualize correlation within region and within/across LD bins; it annotates variant positions according to the LD-bin assignment.

## 3 Usage case

In **<u>Supplementary Information 3</u>**, we demonstrate RegionScan capabilities and computational efficiency by genome-wide analysis of 1,340 individuals with type 1 diabetes (T1D) from the DCCT/EDIC^22,23^ genetic study for LDL-cholesterol (LDL-c, measured at baseline). **Fig. 1** gives an overview of RegionScan capabilities based on the results for chr19. In **<u>Supplementary Information 4</u>**, we report computation time by sample size and region size based on a realistic test dataset.

**Fig. 1.**
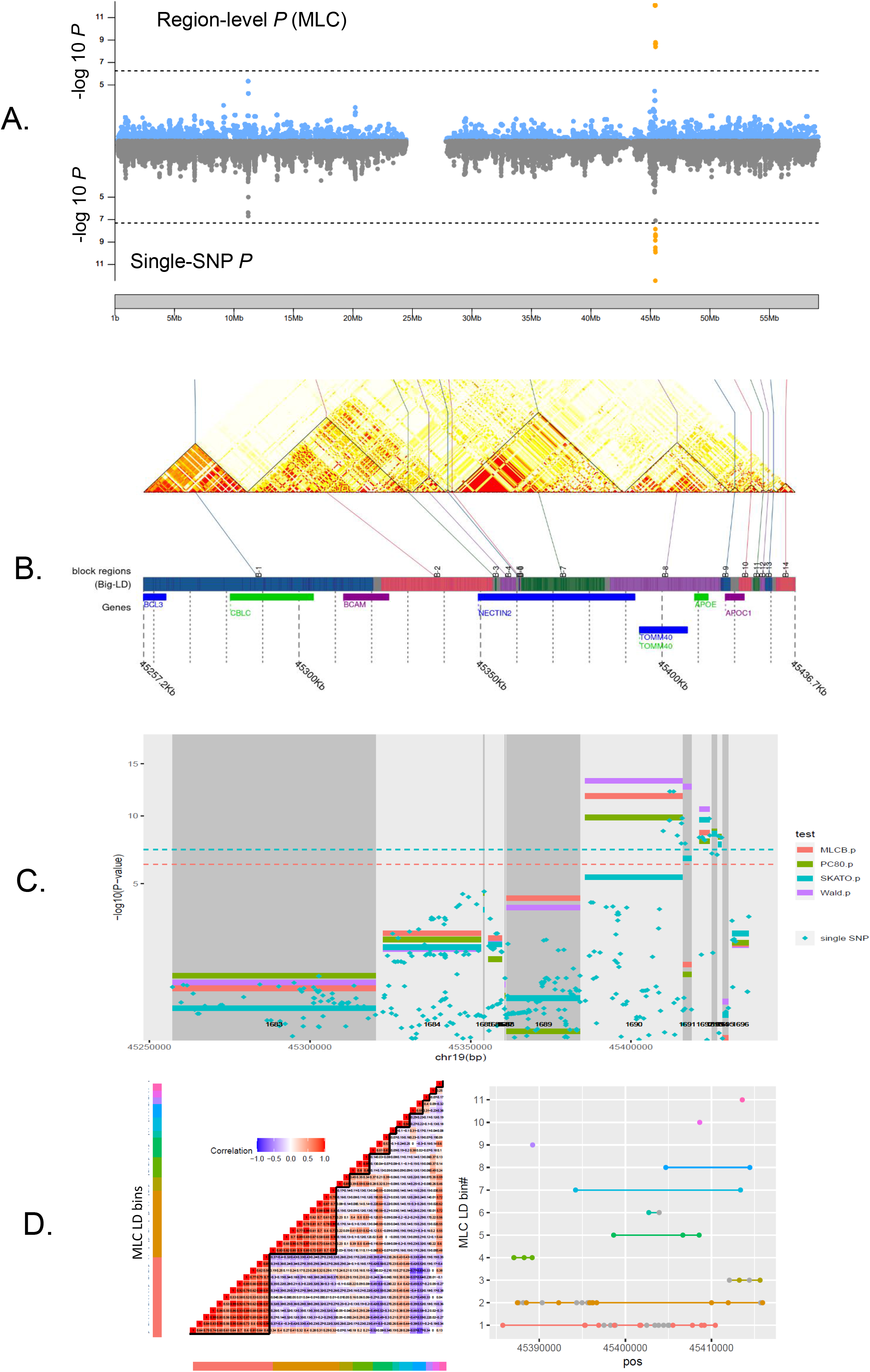
Overview of *RegionScan* for genome-wide region-level association analysis of LDL-c at baseline in 1,340 individuals from the DCCT/EDIC^22,23^ Genetics Study of the Usage case study. Details of analysis are described in **<u>Supplementary Information 3</u>**. To facilitate visualization of the results, we illustrate results on chr19 which exhibits genome-wide region-level association signals at the region- and variant-levels. Panel (**A**) illustrates a comparison between region-level association results based, for example, on the MLC test (top panel) with single-SNP results (bottom panel) for 89,001 regions analyzed genome-wide; the dotted lines indicate the genome-wide Bonferroni-corrected significance levels: 5.6E-7 for region-level tests (top panel) and 5E-8 (bottom panel) for single-SNP tests. Panel (**B**) illustrates partitioning results for 13 regions in chr19: 45,257,201-45,436,657bp; gene positions are shown in GRCh37. The blocks are delimited by triangles. Panel (**C**) illustrates comparison of results for multiple region-level tests for the same 13 regions as illustrated in Panel (**B**); changes in grey shading facilitate visualization of region boundaries. Panel (**D**) shows the LD bins constructed for the MLC test within the top LDL-c associated region, overlapping *APOE* gene (chr19: region #1690, chr19:45,385,759-45,415,935). The left panel shows the heatmap of the SNP correlation matrix (with SNPs ordered by position within LD bin, and LD bins ordered by number of SNPs assigned); the right panel shows the SNP positions (*X* axis) along the LD bins (*Y* axis).

## Conclusion

*RegionScan* is a flexible and versatile R package designed for scalable and comprehensive genome-wide region-level analysis that leverages region definition adaptive to local LD structure (or any other user-provided region definition). It implements multiple region-level tests sensitive to heterogeneous genetic architecture, including LD-bin reduced-*df* region-level tests, facilitates comparisons of region-level and single-variant test results, and includes options to deal with high dimensionality and multi-collinearity arising from improving resolution of genotyped/sequenced/imputed genetic data. Modular design is flexible for future developments.

## Funding

This project was supported by: CIHR Project Grant (#PJT-159463), CIHR STAGE fellowship (MB #GET-101831) and NSERC Discovery Grant (SBB #RGPIN-05896). **Conflict of Interest:** none declared.

## Software, code and data availability

The R package RegionScan (https://github.com/brossardMyriam/RegionScan) is available on GitHub and includes a vignette on how to install and run RegionScan in a realistic artificial dataset provided. The DCCT/EDIC data are available to authorized users at https://repository.niddk.nih.gov/studies/edic/ and https://www.ncbi.nlm.nih.gov/projects/gap/cgi-bin/study.cgi?study_id=phs000086.v3.p1 (IRB #07-0208-E). Data analysis, software development, and computation time estimation were performed on the Hospital for Sick Children High-performance Computing Facility, the Lunenfeld-Tanenbaum Research Institute High-performance Computing platform, and the Niagara supercomputer (with support from the Canada Foundation for Innovation under the auspices of Compute Canada, the Government of Ontario, Ontario Research Fund - Research Excellence, and the University of Toronto). For hardware specifications on Niagara, see https://docs.computecanada.ca/wiki/Niagara#Niagara_hardware_specifications and https://docs.scinet.utoronto.ca/index.php/Niagara_Quickstart.

## Acknowledgements

This study uses data provided by the Diabetes Control and Complications Trial / Epidemiology of Diabetes Interventions and Complications (DCCT/EDIC) Research Group which is sponsored through research contracts from the National Institute of Diabetes, Endocrinology and Metabolic Diseases of the National Institute of Diabetes and Digestive Kidney Diseases (NIDDK) and the National Institutes of Health (NIH). The authors are grateful to the subjects in the DCCT/EDIC cohort for their long-term participation. A complete list of the individuals and institutions participating in the DCCT/EDIC Research Group can be found in **<u>Supplementary Information 3</u>**. This project was supported by: CIHR Project Grant (#PJT-159463), CIHR STAGE fellowship (MB #GET-101831).

## Notes

https://github.com/brossardMyriam/RegionScan

### Competing Interest Statement

The authors have declared no competing interest.

