## Supplementary Information for "RegionScan: A comprehensive R package for region-level genome-wide association testing with integration and visualization of multiple-variant and single-variant hypothesis testing"

### Supplementary Information 1. Region-level tests implemented in the RegionScan R package

In this section, we assume a region with *K* SNPs genotyped/imputed in *N* individuals, and define:

- ***G***, a *N*-by-*K* genotypes matrix;
- ***Y,*** a *N*-by-1 phenotypic vector (qualitative or binary),
- $\boldsymbol{X}$, a matrix of covariates (if applicable)

We classify the region-level tests implemented in RegionScan into three categories: multi-SNP linear/logistic regression-based tests, variance component score tests, and region-level minimum *P*-like tests (**Table S1**).

***Regression-based tests***: For the category of multiple-SNP linear/logistic regression tests, we assume a regional multi-SNP regression model:

$g\left( E[\boldsymbol{Y}] \right)=\beta_{0}+\boldsymbol{G\beta+X\alpha}$	(Equation 1)

Where:

- $g\left( . \right)$ is a link function and can be set to the identity function, if ***Y*** is quantitative or to the logit function if ***Y*** is binary,
- $\boldsymbol{\beta}={(\beta_{1},\ldots,\beta_{k},\ldots,\beta_{K})}^{T}$ is the *K*-length vector of SNP effects,
- $\boldsymbol{\alpha}$ is a vector of covariate effects (if applicable).

The multi-SNP linear/logistic regression-based tests, Generalized Wald test, LC and MLC tests, described in Table 1.1 are all based on the SNP effect estimates vector $\hat{\boldsymbol{\beta}}$ and their variance-covariance matrix $\boldsymbol{\Sigma}$ estimated from the regression model defined in Equation 1.

***Variance component score tests:*** The region-level tests are defined by a statistic $Q={(\boldsymbol{Y}-\hat{\boldsymbol{\mu}_{\boldsymbol{0}}})}^{T}\boldsymbol{H}(\boldsymbol{Y}-\hat{\boldsymbol{\mu}_{\boldsymbol{0}}})$ with $\hat{\boldsymbol{\mu}_{\boldsymbol{0}}}=$predicted phenotypes obtained by fitting the following null genetic model (without SNP): $g\left( E[\boldsymbol{Y}] \right)=\beta_{0}+\boldsymbol{X\alpha}$, $\boldsymbol{H=GWW}\boldsymbol{G}^{T}$ a *N*-by-*N* kernel matrix (positive semidefinite) and $\boldsymbol{W=}diag(w_{1},...,w_{K})$, a matrix of specified SNP weights.

***minP-like tests***: For the region-level min*P*-like tests, we define $\boldsymbol{Z}\mathbf{=}(z_{1},\ldots,z_{k},\ldots,z_{K})$, as the vector of Z scores from single-SNP regression models applied to each of the *K* SNPs ($\boldsymbol{G}_{\boldsymbol{k}}, 1\leq k\leq K)$ in the region, that is:

$$g\left( E[\boldsymbol{Y}] \right)=\beta_{0k}+G_{k}\beta_{k}\boldsymbol{+X}\boldsymbol{\alpha}_{\boldsymbol{k}}$$

We define $p_{k}$as the *P*-value from the 1 degree of freedom (*df*) Wald test for each SNP effect $\beta_{k} ($H0: $\beta_{k}=0$ *vs* H1: $\beta_{k}\neq0)$, and denote $p_{(1)}\leq\ldots\leq$ $p_{(k)}\ldots\leq$ $p_{(K)}$ as the single-SNP *P*-values ranked by increasing order*.*

The Generalized Wald test, LC, MLC, PC80, SKAT, SKAT-O and Min-*P* region-level tests from **Table S1** have been previously compared in simulation studies under various scenarios of complex genetic architectures (see Yoo et *al*^1^ for details).

In **Supplementary Information 2**, we describe steps involved in variant processing and analysis, as well as options implemented in the main *regscan* function to help diagnose and reduce multicollinearity in the regional multi-SNP regression models.

**Table S1.** Features and options of the region-level tests implemented in RegionScan

| **Class of region-level test** | **Region-level tests** | **Method/Features** | **Test Statistic** | **Default parameter values in *regscan*** |
| --- | --- | --- | --- | --- |
| multi-SNP linear/logistic regression tests^a^ | Wald  (default) | Generalized Wald test  (Quadratic test, multi-directional) | $G_{Wald}={\hat{\boldsymbol{\beta}}}^{\boldsymbol{T}}\boldsymbol{\Sigma}^{\boldsymbol{-}\boldsymbol{1}}\hat{\boldsymbol{\beta}}$  H0: $\beta_{k}=$0 **for *all* *K* SNPs** *vs* H1: ***at least* one** $\beta_{k}$ ≠ 0  which is a ***multi-directional* alternative**  Under H0: $G_{Wald}\sim\chi^{2}$with *K df* | None |
|  | LC^2–4^  (default) | Linear Combination test  (Linear test, uni-directional)  The *K* SNPs in region are recoded to maximize the number of SNP pairs with **positive correlation**  $G_{LC}$ power is optimized when SNPs have the same direction of effects | $G_{LC}=$ $\left( \boldsymbol{C}^{\boldsymbol{T}}\hat{\boldsymbol{\beta}} \right)^{T}\left( \boldsymbol{C}^{\boldsymbol{T}}\boldsymbol{\Sigma} \boldsymbol{C} \right)^{-1}\left( \boldsymbol{C}^{\boldsymbol{T}}\hat{\boldsymbol{\beta}} \right)$  Where:   - $\boldsymbol{C}$: (*K-*by*-*1) linear contrast vector,   with $\boldsymbol{C}=\left( \boldsymbol{\Sigma}^{\boldsymbol{-1}}\boldsymbol{J} \right)\left( \boldsymbol{J}^{\boldsymbol{T}}\boldsymbol{\Sigma}^{\boldsymbol{-1}}\boldsymbol{J} \right)^{-1}$ 🡪 $\boldsymbol{C}^{\boldsymbol{T}}$ = $\left( c_{1},\ldots,c_{k},\ldots c_{K} \right)$   - ***J***: is the (*K-*by-1) vector of 1*’s* that assigns all *K* SNPs to one single bin   For δ = $\boldsymbol{C}^{\boldsymbol{T}}\boldsymbol{\beta}$*,* H0: δ = 0 vs H1: δ ≠ 0  which is a ***uni-directional* alternative**  Under H0: $G_{LC}\sim\chi^{2}$with 1 *df* | None  If *alltests*=TRUE, additional equivalent LC test but based on *Z*-scores (noted LCZ) is provided in output (see Yoo et *al*^1^ for mathematical details) |
|  | MLC^5^  (default) | Multiple Linear Combinations test  Reduced-*df* test adaptive to LD within region  (Hybrid linear – quadratic combination test)  The *K* SNPs are clustered into *L* LD bins of correlated SNPs using the CLQ algorithm^6^; within each LD bin, the SNPs are **recoded** to ***maximize*** the number of SNP pairs with ***positive* correlation.**  Note: MLC includes as special cases, the Generalized Wald test (when *L*=*K*) and the LC test (when *L*=1). | $G_{MLC}=\left( \boldsymbol{W}^{\boldsymbol{T}}\hat{\boldsymbol{\beta}} \right)^{T}\left( \boldsymbol{W}^{\boldsymbol{T}}\boldsymbol{\Sigma} \boldsymbol{W} \right)^{-1}\left( \boldsymbol{W}^{\boldsymbol{T}}\hat{\boldsymbol{\beta}} \right)$  Where:   - $\boldsymbol{\beta}^{T}$**= (**$\boldsymbol{\beta}$***_1_^T^ \|* … \|** $\boldsymbol{\beta}$***_l_^T^* \| … \|** $\boldsymbol{\beta}$***_L_^T^* ) &** $\boldsymbol{\Sigma}$, are ordered by LD bin, such that: $\boldsymbol{\beta}_{\boldsymbol{l}}\boldsymbol{=(}\beta_{k_{l,1}},\ldots,\beta_{k_{l,1}},\ldots,\beta_{K_{l}})$ corresponds to the vector of SNP effects for all $K_{l}$ SNPs assigned to the *l*^th^ LD bin. - $\boldsymbol{W}$ $=\left( \boldsymbol{\Sigma}^{\boldsymbol{-1}}\boldsymbol{J} \right)\left( \boldsymbol{J}^{\boldsymbol{T}}\boldsymbol{\Sigma}^{\boldsymbol{-1}}\boldsymbol{J} \right)^{-1}$is a (*K-*by*-L*) matrix of linear contrasts - ***J* = [ *J_1_ \| J_2_* \| … \| *J_L_* ]** is the (*K-*by*-L)* indicator *matrix* that assigns *K* SNPs to *L* LD bins 🡪 ***W^T^* = [ *W_1_ \| W_2_* \| … \| *W_L_* ]*^T^***, where ***W_l_^T^*** = [ $\boldsymbol{W}$***_l1_^T^ \|*** $\boldsymbol{W}$***_l2_^T^* \| … \|** $\boldsymbol{W}$***_ll_^T^* \| … \|** $\boldsymbol{W}$***_lL_^T^*** ] is (1 by *K*) - $\boldsymbol{W}$***_ll_*** and $\boldsymbol{W}$***_lm_***: LD *within* and *between* LD bins   For $\boldsymbol{\delta}=\boldsymbol{W}^{\boldsymbol{T}}\boldsymbol{\beta}$*,* where each δ*_l_* = $\boldsymbol{W}$***_l_^T^*** $\boldsymbol{\beta}$ corresponds to the bin-specific effect for *l*^th^ bin  H0: δ*_l_* = 0 **for all *L* LD bins** vs H1: **at least *one*** δ*_l_* ≠ 0,  which is a ***reduced-dimension multi-directional alternative*** (sensitive to directional effects defined by each LD bin)  $\boldsymbol{G}_{\boldsymbol{MLC}}\boldsymbol{\sim}\boldsymbol{\chi}^{\boldsymbol{2}}$with ***L*** $\boldsymbol{\leq}\boldsymbol{K}$ ***df*** | Clustering parameter:  *edgecut*=0.50  (minimum absolute correlation between SNPs in a LD bin, note: corresponds to r2 =0.25)  If *alltests*=TRUE, additional equivalent MLC test but based on *Z*-scores (noted MLCZ) is provided in output (see Yoo et *al*^1^ for mathematical details) |
|  | PC80^7^  (default) | **Dimension reduction step:** Principal component analysis of the *K* SNPs, selection of the *S* PC-SNPs, that explain at least PCcut=80% of the total SNP variance  Multiple regression of the *S* PC-SNPs & Generalized Wald test  Reduced-df test (*S* $\leq$*K*)  (multi-directional) | $G_{PC80}={\hat{\boldsymbol{\gamma}}}^{T}\boldsymbol{\Pi}^{-1}\hat{\boldsymbol{\gamma}}$  $G_{PC80}\sim\chi_{S}^{2}$with *S* $\leq K$ *df*  Where: $\hat{\boldsymbol{\gamma}}=(\hat{\gamma_{1}},\ldots,\hat{\gamma_{s}},\ldots,\hat{\gamma_{S}})$ is the vector of PC-SNPs effects and $\boldsymbol{\Pi}$ is the corresponding variance-covariance matrix,  estimated by multiple regression.  H0: $\gamma_{s}=0$ **for *all* *S* PC-SNPs** vs H1: **at *least* one** $\gamma_{s}\neq0$  which is a ***reduced-dimension multi-directional alternative*** | PCcut: Variation explained by the PCs selected (default PCcut=80%) |
| Variance component score tests | SKAT^8,9^  (default) | Variance-like component test under a logistic/linear mixed model | **Linear model assumed under SKAT**  Variance-component score statistic:  $Q_{SKAT}={(\boldsymbol{Y}-\hat{\boldsymbol{\mu}_{\boldsymbol{0}}})}^{T}\boldsymbol{H}(\boldsymbol{Y}-\hat{\boldsymbol{\mu}_{\boldsymbol{0}}})$  Where:   - $\hat{\boldsymbol{\mu}_{\boldsymbol{0}}}:$ predicted phenotypes obtained by fitting the null genetic model (without SNPs) - $\boldsymbol{H=GWW}\boldsymbol{G}^{T}$ : a *N*-by-*N* kernel matrix (positive semidefinite)   $Q_{SKAT}$can be rewritten as:  $Q_{SKAT}=\sum_{k=1}^{K} w_{k}^{2}\left[ \sum_{i=1}^{N} (y_{i}-\hat{\mu_{0,i}})g_{i,k} \right]$  H0: $\tau=0 (\Longleftrightarrow\boldsymbol{b}=0$)  H1: $\tau>0 (\Longleftrightarrow\boldsymbol{b}\neq0)$ (at least one genetic effect)  Under H0: $Q_{SKAT}$ follows a complicated mixture of $\chi_{1}^{2}$ distributions^10^. | Based on SKAT function R SKAT^11^ R package with following option:  kernel="linear",  No weights used  In “region output”,  columns named:   - “SKAT, SKAT.pLiu and SKAT.pDavies” corresponds to SKAT results with $\boldsymbol{W=}\boldsymbol{I}_{\boldsymbol{K}}$; - *P*-values computed using both: an exact method (“davies) and an approximation method (“liu”) |
|  | SKATO^12^  (default) | Combined test of SKAT and burden test ($Q_{SKAT-B})$  $Q_{SKAT-B}$ power is optimized when SNPs have the same direction of effects and high correlation  SKAT-O includes SKAT ($\rho=1)$ and burden test as special cases ($\rho=0)$ | $Q_{SKAT-O}={\rho Q}_{SKAT}+(1-\rho)Q_{SKAT-B}$  Where:   - $0\leq\rho\leq1$, an optimal $\rho$ is computed from a grid search to maximise the power^12^ - $Q_{SKAT-B}=\left[ \sum_{i=1}^{N} (y_{i}-\hat{\mu_{0,i}})\left( \sum_{k=1}^{K} w_{k}g_{i,k} \right) \right]^{2}$   Under H0: $Q_{SKATO} \sim$mixture of $\chi_{1}^{2}$ distributions (as for the SKAT statistic) $+ \chi^{2}(1df)$ | Based on SKAT function from “SKAT^11^” R package with following options:  kernel="linear.weighted",  weights.beta=c($a_{1},a_{2}$),  method="optimal.adj"  $a_{1}=a_{2}=0.5$corresponds to weights used by Madsen and Browning^13^ for set-based analysis of common variants^14^    "optimal.adj" computes $Q_{SKAT-O}$ P-values based on a grid search of *ρ*) and then uses the minimum *P*-value as a test statistic, as previously recommended^15^ |
| region-level min*P-like* tests^a^ | MinPJ^16^  (if *alltests*=  TRUE) | Minimum single-SNP P-value in the region, corrected for the number of tests  Note: *can be computationally expensive due to fitting a generalized estimating equation model (GEE) to estimate the variance-covariance matrix (*$\boldsymbol{\Lambda}$*)* | Let $\boldsymbol{Z=}\left( Z_{1},\ldots,Z_{k},\ldots,Z_{K} \right)$ the vector of Z scores from single-SNP analysis and $\boldsymbol{\Lambda}$, their covariance matrix estimated using the robust GEE variance method as previously suggested^17^.  $\boldsymbol{Z}\sim N_{K}(\boldsymbol{0}, \boldsymbol{\Lambda})$,  $Min P=1-P\left( max(\boldsymbol{Z})<\Phi^{-1}\left( 1-\frac{p_{(1)}}{2} \right) \right)$  Computation of $Min P$ requires integration of the multivariate normal density function, which has not a closed-form solution. Integration possible with mtvnorm R package^18^ but can be computationally intensive for the analysis of large regions. In RegionScan, we implemented the approximation of the normal probabilities proposed by James et *al*^19^ to improve computational efficiency for genome-wide region-level analysis. | None |
|  | GATES^20^  (default) | Minimum single-SNP *P*-value corrected for the effective # of SNPs tested in the region using the Extended Simes procedure. | $P_{GATES}=min\left( \frac{m\times p_{(j)}}{m_{(j)}} \right)$  Where:   - $m$: effective number of independent *P*-values in the region, with: $m=M-\sum_{j=1}^{M} \left[ I(\lambda_{j}>1)(\lambda_{j}-1) \right]\lambda_{j}$, where *I(*x) is an indicator function and $\lambda_{j}$ are the eigenvalues of the correlation matrix of the single-SNP *P*-values. This correlation matrix is approximated by a sixth-order polynomial function of the pair-wise allelic correlation coefficient of the regional SNP matrix^20^ - $m_{j}:$effective number of independent *P*-values in the region, calculated as *m* but for the top *j* *P*-values in the region | None |
|  | SimpleM^21^  (default) | Minimum single-SNP *P*-value **corrected for the effective # of SNPs** tested in the region & their LD | $P_{SimpleM}=1-{(1-p_{\left( 1 \right)})}^{m}$  Where: $m$= number of eigenvalues from the SNP correlation matrix of the *K* SNPs ***ranked*** by descending order, that explain cumulatively at least SimpleM_cut=0.995 of the SNP variation in the region | *SimpleMcut*=0.995 |
|  | uMinP  (default) | Minimum single-SNP *P*-value **uncorrected**  (reported to facilitate comparison with single-SNP results) | $P_{uMinpP}=p_{(1)}$ | None |

^a^SNP effects and their variance-covariances estimated using the generalized “glm” function from the R “stats” package or, when *firthreg* =TRUE from the “brglm2” R package.

### Supplementary Information 2. Details on the main *regscan* function and auxiliary *recodeVCF* function

#### 2.1 *regscan*: main function to process and analyze regions

##### 2.1.1 Steps involved in processing, and analysis of each region

In **Fig. S1,** we summarize the main steps implemented in *regscan* from data processing to analysis of each region, with details described thereafter.

**Fig. S1.** Overview of the steps in *regscan*

Step 1: SNP recoding

recoding of all SNPs such as baseline allele = major allele

Step 2: SNP filtering/selection

filtering of the SNPs with MAF < mafcut (default mafcut= 0.05)

Step 3: Clustering & Recoding (for the MLC test)

Clustering of the bialleic SNPs into LD bins (default edgecut=0.5)

Recoding of the biallelic SNPs to maximize the # of SNP pairs with positive correlation within each LD bin

If multiallelic = TRUE, assignation of the multiallelic-SNPs to LD bins based on correlation

Step 4: SNP pruning

Pruning of the SNPs in complete linear dependency (aliasing) to reduce fitting issues in multiple-SNP regional regression models

Step 5: Region-level & SNP-level analysis

Region-level & single-SNP tests applied to all K remaining SNPs after Step 4

if singleSNPall=TRUE: report single-SNP test results applied to all the SNPs kept at Step 2 (this includes all the SNPs before pruning on LD/complete linear dependency)

***Step 1. Recoding of all the SNPs such as baseline allele = major allele***

In *regscan,* all the SNPs within each region are automatically recoded such that their baseline allele = major allele, as some of the region-level tests are sensitive to the direction of SNP effects (**Supplementary Information 1**). When the analysis is based on the bi-allelic SNPs only (multiallelic=FALSE), *regscan* automatically recodes all the SNPs such that their baseline alleles match the major allele. When the analysis also includes multi-allelic SNPs (multiallelic=TRUE), then a VCF file name must be specified and *regscan* will use internally *recodeVCF* to automatically create *geno/SNPinfo* inputs for each region and apply an algorithm we implemented to recode the multi-allelic SNPs such that the baseline matches the major allele (see details in **section 2.2**).

***Step 2. SNP filtering/selection***

For region-level regression tests such as implemented in RegionScan , SNPs with low MAFin regions can lead to numerical instability in model fitting. By default, *regscan* pruns out the SNPs with MAF < 0.05 (Step 1). For studies with large sample sizes, the default *mafcut* value can be relaxed to select a subset of SNPs based on a MAF criteria (or low minor allele count, MAC).

***Step 3. Clustering and recoding of the SNPs for the MLC region-level test***

i. **Clustering** **of the biallelic SNPs** **in LD bins** based on the Pearson correlation coefficient metric (see Yoo et *al*^6^ for details on the CliQue-Based (CLQ) clustering algorithm implemented in RegionScan). By default, the clustering parameter (*edgecut*) is set to 0.5 based on recommendations from numerical experimentations^1^. The default parameter value edgecut=0.5, corresponds to LD as defined by r^2^ = 0.25.
ii. **Recoding of the biallelic SNPs** within each LD bin to maximise the no. of biallelic SNP pairs with **positive correlation** for the MLC test.
iii. **if *multiallelic* = TRUE: assign each multi-allelic SNP to an LD bin** based on correlation:
     a) Identification of the bin(s) with minimum correlation with the multi-allelic SNP variable ≥ *edgecut* & assign to the LD bin **with the largest mean correlation** with that multi-allelic SNP variable
     b) Otherwise, if **no bin matches the condition in a)**: assign the multi-allelic SNP variable to a new LD bin & update the list of LD bins
     c) Repeat the previous steps for all remaining multi-allelic SNP variables (note: each alternate allele of the same multi-allelic SNP can be assigned to different LD bins)

***Step 4. Pruning of highly correlated SNPs & SNPs in complete linear dependency***

Perfect correlation/high multicollinearity in regions can lead to the inability to estimate the multiple-SNP regression models (e.g. ill-conditional regional genotypes matrix). Therefore, by default in *regscan,* within each region we prune out SNPs with Pearson correlation (in absolute value) larger than 0.99 (*rcut*). The *rcut* value can be adjusted, for example based on the study sample size, and type of trait studied (quantitative or binary). LD pruning is skipped if LDpruning=FALSE. In the latter case, aliases will be checked before fitting the multi-SNP regional regression model and removed from sets of analyzed SNPs for the region-level tests. In the variant-output level file, we also report VIF values to diagnose multicollinearity in multi-SNP regression models at the LD bin or region levels.

1. **Pruning of the SNPs *highly* correlated within each LD bin** (default *rcut*=0.99):

If *multiallelic* *=* FALSE: pruning of the bi-allelic SNPs

If *multiallelic* *=* TRUE

a. For the multi-allelic SNPs: Distinguish two sets of multi-allelic SNPs:

setA: multi-allelic SNPs with *only one* bi-allelic SNP variable left

setB: multi-allelic SNPs with at *least two* bi-allelic SNP variables left

b. Then, pruning of the SNPs on the following order:

Pruning of the bi-allelic SNPs,

Pruning of the setA SNPs

If a bi-allelic SNP has abs(r)≥*rcut* with a SNP from setA;

keep the bi-allelic SNP and exclude the setA SNP.

Pruning of the setB SNPs

2. **Identification of aliases within the region** using the *alias* function from the base R “stats” package.

***Step 5. Region-level tests & Single-SNP tests***

All the region-level tests implemented in *regscan* are based on the exact same set of *K* SNPs kept after **Step 4** to facilitate comparisons of the results across the different tests. For the LC test, the bi-allelic SNPs within each region are recoded in *regscan* to maximize the number of SNP pairs with ***positive* correlation** within the region. The variant-level output includes indicator variables to identify the SNPs that have been recoded for MLC (MLC.flip) or LC (LC.flip) region-level tests (see **section 2.1.3**). The single-SNP analysis is based on all the SNPs kept after **Step 1** (SNPs unpruned on correlation or complete linear dependency).

**Fig. S2.** Illustration of the SNP clustering and recoding in step2 for the MLC region-level test. These plots were produced using the option MLCheatmap=TRUE in the *regsan* main function

(A.) Heatmap of the SNP correlation matrix within the region ***before* pruning, clustering and recoding** of the SNPs; with SNPs ordered by physical position on chr 19

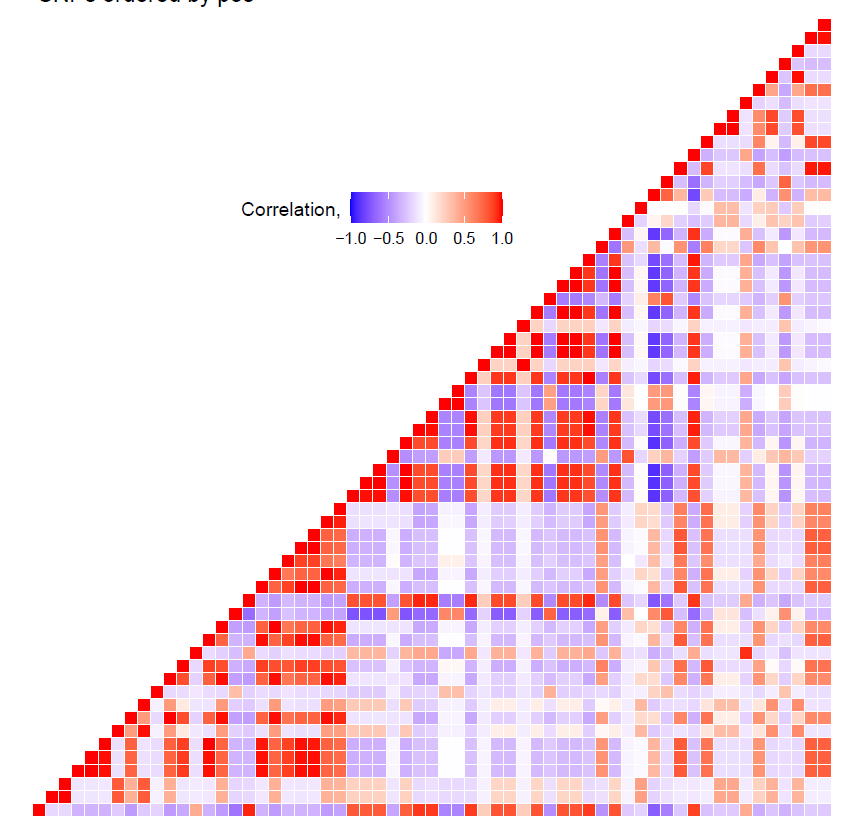

(B.) Heatmap of the SNP correlation matrix within the region **after SNP clustering and before pruning & recording**; with **SNPs ordered by LD** (from the largest LD bin to the smallest bin) and by position within each LD bin. Each LD bin is assigned a different color for visualization and and match panel (D.)

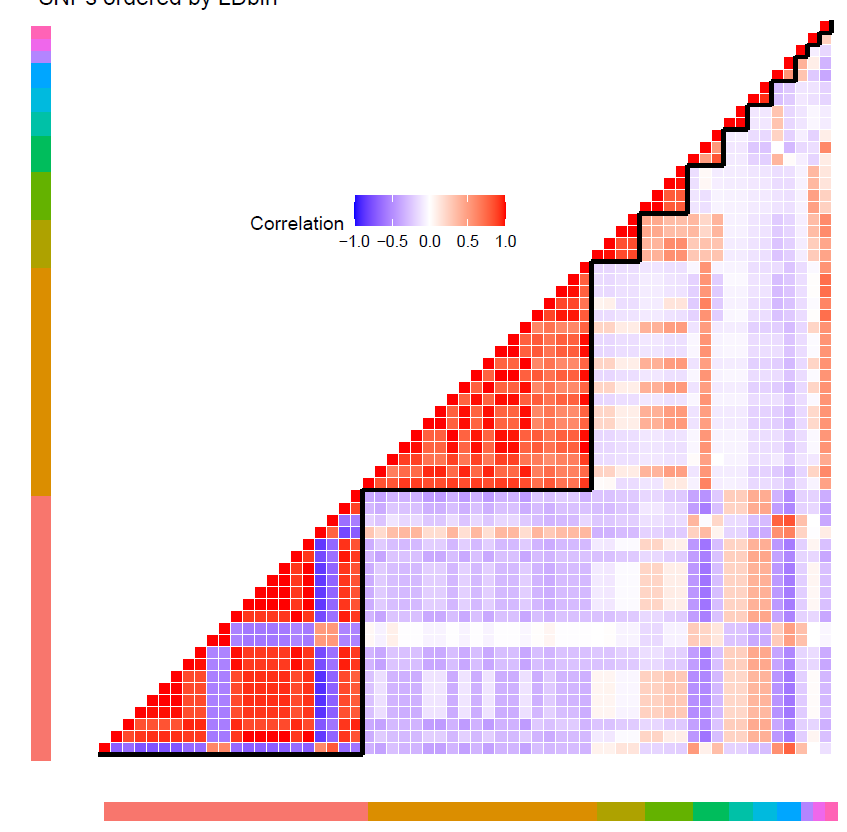

(C.) Heatmap of the SNP correlation matrix within the region **after SNP clustering & pruning but before recording**; with **SNPs ordered by LD** (from the largest LD bin to the smallest bin) and by position within each LD bin.

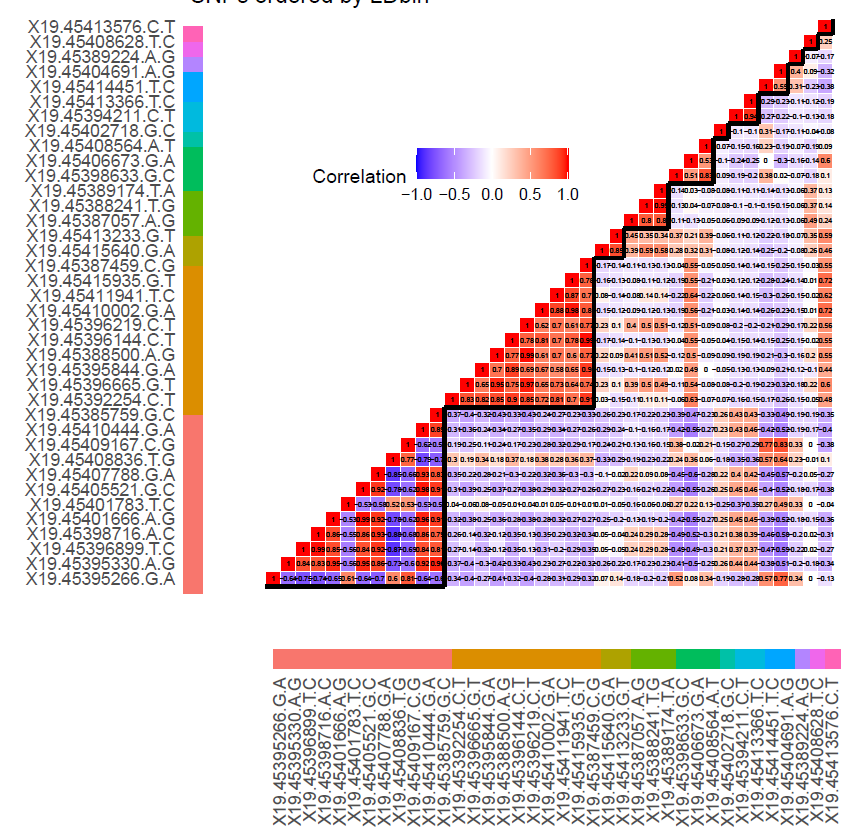

(D.) Heatmap of the SNP correlation matrix within the region as for panel (B.) **after clustering, pruning and recoding** within each LD bin to maximize the no. of SNP pairs with positive correlation within LD bin.

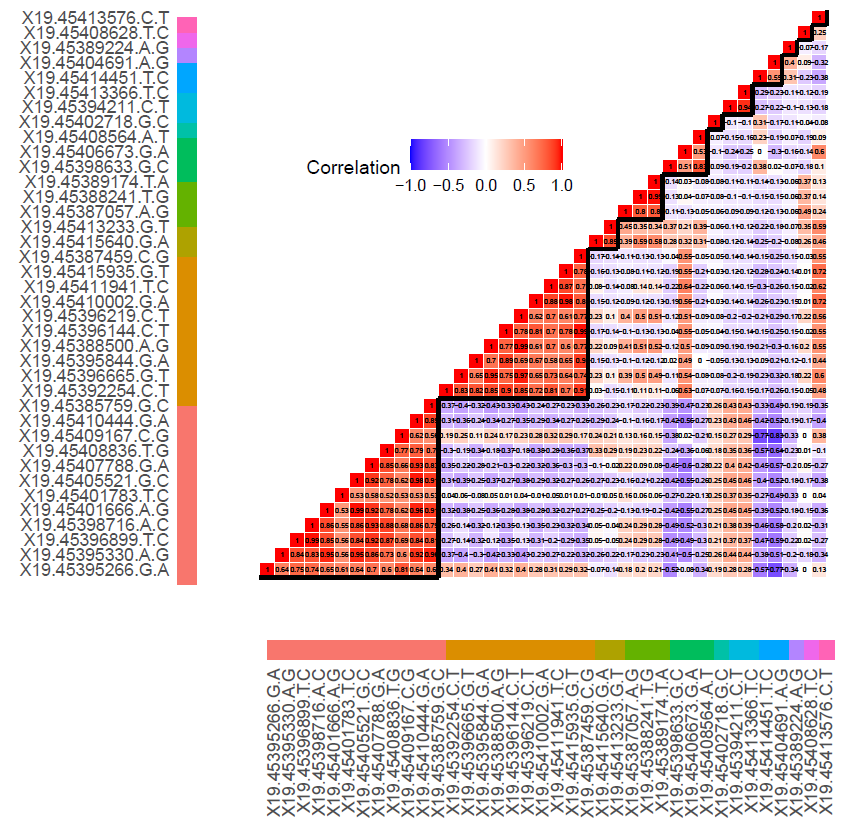

##### 2.1.2 List of main arguments and default values

| **Arguments** | **Definition** | **Default value** |
| --- | --- | --- |
| REGIONinfo | dataframe including region definition, with at least 4 columns: “chr”, “region”, “start.bp”, “end.bp”, as provided for example, by the output of “BigLD” function from the *gpart* R package | Required |
| phenocov | dataframe including the covariates (if applicable) and the phenotype (s) in columns. Individuals (in rows) must be in the same order as in the data file. |  |
| covlist | list of covariates to account for in single-SNP and multiple-SNP regression models, including for example, ancestry PCs and/or non-genetic covariates, to include in the analysis (must be included in input *data* or *phenocov* if it applies) | NULL |
| covout | Option to output the covariates coefficients, standard errors and their P-values from multi-SNP regional regression (used to construct MLC, LC and Generalized Wald tests) | FALSE |
| pheno | Name of the phenotype column in *phenocov* | Required |
| pheno_type | pheno_type=“C” if phenotype is **c**ontinuous,  or pheno_type=“D” if phenotype is **d**ichotomous | Required |
| geno_type | Format of the genotypes: geno_type=“D” if genotypes are in allele dosage format , or genotype format (geno_type=“G”)  – argument required for SKAT/SKATO tests | Required |
| ***If geno/info data frames are specified* (in this case, recodeVCF is not used)** | |  |
| data | Dataframe including the genotypes in columns | Required if *vcfname* is left empty |
| SNPinfo | Dataframe including the SNP information,  must include the following columns:  "chr", "pos", "variant", "ref", "alt", “multialSNP”, "multialSNP.rec","freq.alt"   - "multialSNP": binary variable=0, if the SNP is bi-allelic (two alleles), or 1 if more than two alleles - "multialSNP.rec": indicates if the multi-allelic SNP has been recoded b recodeVCF (such that baseline allele = major allele) - “freq.alt": frequency of the alternate allele (minor allele)   This input can be generated using the auxiliary function *recodeVCF* | Required if *vcfname* is left empty |
| ***Used only by the auxiliary function recodeVCF***  ***if data and SNPinfo are not specified*** | |  |
| vcfname | name of the vcf file (if required) | NULL |
| qcmachr2 | threshold used to filter out SNPs with low Mach R2 imputation quality score | NULL |
| qcinput | Dataframe including at least 2 columns: variant & info_score (imputation quality score) | NULL |
| info_score | threshold used to filter out SNPs with low Info imputation quality score – If this option is specified, “qcinput” must be specified, and “info_score” must be a column in “qcinput” | NULL |
| ***Options to include multiallelic SNPs*** | |  |
| multiallelic | If FALSE, extract & process only the biallelic SNPs,  If TRUE: include multiallelic SNPs in addition of the biallelic SNPs | FALSE |
| multial_nmaxalleles | For extraction of SNPs with less than multial_nmaxalleles alleles (e.g. 2 to extract biallelic SNPs (default), 3 to extract bi-allelic and triallelic SNPs, etc) | 2 |
| ***Options for pruning/filtering of the SNPs in each region***  ***(see details in Supplementary Information 2, section 2.3)*** | |  |
| mafcut | Threshold to filter out SNPs with MAF below threshold | 0.05 |
| LDpruning | Option to prune out SNPs within region based on absolute value of the regional correlation matrix; recommended to reduce multi-collinearity issues in multi-SNP regional regression models (see section 2.1.2). | TRUE |
| rcut | Works only if LDpruning=TRUE, threshold to prune out SNPs based on correlation (in absolute value) | 0.99 |
| ***Other options*** | |  |
| singleSNPall | Produces an additional output including single-SNP results (and LD-bin level information) for all SNPs before LD pruning/alias identification.  – can increase computational time. | FALSE |
| firthreg | Works only if pheno_type=“D”: If firthreg =“TRUE”, use a Jeffreys-prior penalized-likelihood regression implemented in the R package **brlmgm2**^22,23^ instead of the default maximum-likelihood logistic regression. This option is recommended for unbalanced case-control data / small minor allele count | FALSE |
| parallel | By default, process & run region-level analysis sequentially. If “TRUE” specified, proceed in parallel | FALSE |
| regionlist | list of the region names from REGIONinfo input to be analyzed (can be a subset of the regions) | NULL |
| alltests | By default, output region-level tests: Wald, MLCB, PC80, SKAT, SKATO, LCB, GATES, SimpleM;  If alltests=“TRUE” reports additional region-level tests: MinPJ, MLCZ, LCZ | FALSE |
| edgecut | parameter for clustering of the SNPs in LD bins (based on SNP correlation) using the CLQ algorithm applied before MLC test | 0.5^a^ |
| tol | Tolerance parameter to deal with convergence issues in regression models | 1e-16 |
| MLCheatmap | For each region, produces four heatmap plots of the regional SNP correlation matrix, with SNPs ordered by positions & by LD bin, before and after LD pruning (if applicable). Option recommended for a subset of regions of interest specified with the option regionlist. | FALSE |

^a^By default, edgecut=0.50 (which corresponds to r2=0.25), showed to be the optimal clustering parameter value by simulation studies.

##### 2.1.3 Description of the main outputs of *regscan* function

**Table S2.** Region-level output includes region positions, and region-level test results for all regions analyzed.

| **Column name** | **Description** |
| --- | --- |
| chr | chromosome # (as provided in *REGIONinfo* input*)* |
| region | region # or name (as provided in *REGIONinfo* input*)* |
| start.bp | start region position in bp (as provided in *REGIONinfo* input) |
| end.bp | end region position in bp (as provided in *REGIONinfo* input) |
| max.VIF | Maximum VIF within region |
| NSNPs | NSNPs in region (before pruning on LD/perfect linear dependency) |
| NSNPs.kept | NSNPs analyzed in region (after pruning on LD/perfect linear dependency) |
| Wald | Generalized Wald statistic |
| Wald.df | Degree of freedom of the Generalized Wald statistic (= NSNPs.kept) |
| MLCB | MLC test statistic (based on SNP effects) |
| MLCB.df | Degree of freedom of the MLCB test ( = # of LD bins in the region) |
| MLCB.p | *P*-value for MLCB test |
| LCB | LCB statistic |
| LCB.df | Degree of freedom of the LCB test (=1 for all regions) |
| LCB.p | *P*-values for LCB test |
| PC80 | PC80 statistic |
| PC80.df | Degree of freedom of the PC80 test |
| PC80.p | *P*-value for PC80 test |
| SKAT | SKAT statistic |
| SKAT.pDavies | *P*-values from SKAT computed using Davie’s method |
| SKAT.pLiu | *P*-values from SKAT computed using Liu’s method |
| SKATO.p | SKATO statistic |
| GATES.df | GATES degree of freedom |
| GATES.p | GATES *P*-value |
| SimpleM.p | simpleM *P*-value |
| uMinP.p | Minimum *P*-value (uncorrected) |
| **If alltests=TRUE** | |
| MinPJ.p | Minimum *P*-value (corrected, see **Table S1**) |
| MLCZ | MLC statistic based on Z-scores from multi-SNP regional regression model  (rather than on SNP effects as used for MLCB) |
| MLCZ.p | *P*-value for MLCZ test statistic from multi-SNP regional regression model  (rather than on SNP effects as used for MLCB) |
| LCZ | LC statistic based on Z-scores (rather than SNP effects) from multi-SNP regional regression model  (rather than on SNP effects as used for MLCB) |
| LCZ.p | *P*-value for LCZ test statistic |

**Table S3.** Bin-level output, includes LDbin specific results within regions analyzed

| **Column name** | **Description** |
| --- | --- |
| chr | chromosome # (as provided in input *REGIONinfo)* |
| region | region # or name (as provided in input *REGIONinfo)* |
| start.bp | start region position in bp (as provided in input *REGIONinfo)* |
| end.bp | end region position in bp (as provided in input *REGIONinfo)* |
| bin | LDbin # assigned within each region (numbered by decreasing # of SNPs) |
| bin.size | # of SNPs in LD bin |
| bin.size.keptSNPs | # of SNPs in LD bin (after pruning on MAF & LD (if applicable),  and complete linear dependency) |
| deltabinB | Bin-level effect |
| deltabinB.p | *P*-value for 1 df Wald test of the bin-level effect |
| **If alltests=TRUE** |  |
| deltabinZ | Same as deltabinB but using Z statistics instead of Betas |
| deltabinZ.p | Same as deltabinB but using Z statistics instead of Betas |

**Table S4.** Variant-level output including variant information/positions as well as LD bin assignation and estimates from single-SNP and multiple-SNP regression models applied within each region and each LD bin for all the SNPs analyzed in region-level tests (kept after pruning on MAF and LD/linear dependency, if applicable)

| **Column name** | **Description** |
| --- | --- |
| chr | chromosome # (as provided in *REGIONinfo* input) |
| region | region # or name (as provided in *REGIONinfo* input) |
| start.bp | start region position in bp (as provided in *REGIONinfo* input) |
| end.bp | end region position in bp (as provided in *REGIONinfo* input) |
| variant | variant name (as provided in *SNPinfo* input) |
| pos | SNP position (as provided in *SNPinfo* input) |
| multiallelicSNP | indicator variable to flag multiallelic SNPs (as provided in *SNPinfo* input) |
| ref | Reference allele (as specified in *SNPinfo* input) |
| alt | Alternate allele (as specified in *SNPinfo* input) |
| maf | Minor allele frequency |
| LDbin | LDbin # assigned within each region (numbered by decreasing # of SNPs) |
| LDbin.size | # of SNPs analyzed in each LDbin (kept after pruning on MAF & LD) |
| MLC.flip | Flag the SNPs recoded for MLC/LCbin tests |
| LC.flip | Flag the SNPs recoded for the LCbin tests |
| sg.beta | SNP effect estimate from single-SNP regression models |
| sg.pval | *P*-value for 1 *df* Wald test of SNP effect from single-SNP regression models |
| VIF | Variance inflation factor (VIF) values based on *all SNPs* analyzed in each region |
| glm.beta | SNP effect estimate from the regional multi-SNP regression model |
| glm.pval | *P*-value for 1 *df* Wald test of SNP effect the regional multi-SNP regression model |

**Table S5.** List of SNPs excluded from the region-level tests and reasons for exclusion

| **Column name** | **Description** |
| --- | --- |
| chr | Chromosome # (as provided in input *REGIONinfo)* |
| region | region # (as provided in input *REGIONinfo)* |
| start.bp | end position in bp (as provided in input *REGIONinfo)* |
| end.bp | start position in bp (as provided in input *REGIONinfo)* |
| variant | variant name (as provided in input *REGIONinfo)* |
| multiallelicSNP | indicator variable to flag multiallelic SNPs (as provided in input *REGIONinfo)* |
| pos | SNP position (as provided in input *SNPInfo)* |
| MAF | Minor allele frequency |
| reason | Reason of exclusion:  “mafcut” – MAF < *mafcut* (if applicable)  “rcut” – high correlation (if applicable)  “alias” – complete linear dependency  “multial” – multiallelic SNP (if applicable) |

**Table S6.** Single-SNP results and bin-level information for all SNPs in regions (before LD pruning – optional)

| **Column name** | **Description** |
| --- | --- |
| chr | Chromosome # (as provided in input *REGIONinfo)* |
| region | region # (as provided in input *REGIONinfo)* |
| start.bp | end position in bp (as provided in input *REGIONinfo)* |
| end.bp | start position in bp (as provided in input *REGIONinfo)* |
| variant | variant name (as provided in input *SNPinfo*) |
| pos | SNP position (as provided in input *SNPinfo*) |
| multiallelicSNP | indicator variable to flag multiallelic SNPs (as provided in input *SNPinfo*) |
| major.allele | Major allele (as provided in input *SNPinfo*) |
| minor.allele | Minor allele (as provided in input *SNPinfo*) |
| maf | Minor allele frequency |
| LDbin | Bin #; within each region (numbered by decreasing # of SNPs) |
| sg.beta | SNP effect estimate from single-SNP regression models |
| sg.pval | *P*-value for 1 *df* Wald test of SNP effect from single-SNP regression models |
| rmcorr | Indicator variable to flag SNPs excluded because of LD pruning |

#### 2.2 *recodeVCF*: auxiliary function to extract and process SNPs in VCF files

In this section, we describe the algorithm implemented in *recodeVCF* to extract (or calculate) the allele dosage and recode the bi-allelic (and multi-allelic SNPs, if *multiallelic*=TRUE) from an **unfiltered** VCF type 4 file format that includes all the imputation probabilities for all the SNPs to produce *data* and *SNPinfo* inputs used in *regscan*.

##### 2.2.1 Processing of bi-allelic SNPs (multiallelic = FALSE)

By default, the options in *regscan/recodeVCF* are set to extract the allele dosage of the bi-allelic SNPs from the vcf file, recode such as their baseline allele = major allele and to filter them on MAF > *mafcut*.

##### 2.2.2 Processing of multi-allelic SNPs (multiallelic = TRUE)

We assume that each multi-allelic SNP is coded in the VCF file as a set of bi-allelic SNP variables, with the same genomic position and reference allele; this is as typically provided from imputation reference panels, such as 1000 Genomes (used in our illustration presented in **Supplementary Information 3**). For each individual, imputation programs typically return two imputation probabilities for each of the biallelic SNP variables that comprise the multi-allelic SNP. The two imputation probabilities correspond, respectively, to the probability that individual *i* has one copy of the alternate allele and to the probability that individual *i* has two copies of the alternate allele (see examples in **Tables S7** and **S8**). However, the reference allele does not always match the major allele, depending on the reference alleles from the reference panel used for imputation, and rationalization is needed as described in what follows.

**Table S7.** Illustration of a fictitious tri-allelic SNP (*M*=3 alleles) coded as two bi-allelic SNP variables with the same position and baseline allele A_1,_ but different alternate alleles (A_2_ and A_3_). The allele frequencies of all 3 alleles are assumed to sum to 1 (ie f(A_1_) + f(A_2_) + f(A_3_) = 1)

| Tri-allelic SNP | Chr:position | Alleles Alternate/reference | Frequency of alternate allele |
| --- | --- | --- | --- |
| snp1_A2 | 19:500003 | $A_{2}$/A_1_ | f($A_{2})$ |
| snp1_A3 | 19:500003 | $A_{3}$/A_1_ | f($A_{3})$ |

**Table S8.** Example of calculation of dosage for each alternate allele of the tri-allelic SNP from Table S7 for an individual *i.*

| SNP | Imputation probabilities | probability that individual *i* has | Dosage of alternate allele |
| --- | --- | --- | --- |
| snp1_A2 | $p_{i}^{\boldsymbol{A}_{\boldsymbol{2}}A_{m}}$, $m\epsilon\{A_{1},A_{3}\}$ | ***exactly one*** copy of $A_{1}$  *(ie.* either $A_{1}A_{2}$ or $A_{1}A_{3})$ | $d_{i}\left( \boldsymbol{A}_{\boldsymbol{2}} \right)=p_{i}^{\boldsymbol{A}_{\boldsymbol{2}}A_{m}}+2p_{i}^{\boldsymbol{A}_{\boldsymbol{2}}\boldsymbol{A}_{\boldsymbol{2}}}$ |
|  | $p_{i}^{A_{2}A_{2}}$ | ***two copies*** of $A_{1}$  *(ie.* $A_{1}A_{1})$ |  |
| snp1_A3 | $p_{i}^{\boldsymbol{A}_{\boldsymbol{3}}A_{m}}$, $m\epsilon\{A_{1},A_{2}\}$ | ***exactly one*** copy of $A_{2}$  *(ie.* either $A_{3}A_{1}$ or $A_{3}A_{2})$ | $d_{i}\left( \boldsymbol{A}_{\boldsymbol{3}} \right)=p_{i}^{\boldsymbol{A}_{\boldsymbol{3}}A_{m}}+2p_{i}^{\boldsymbol{A}_{\boldsymbol{3}}\boldsymbol{A}_{\boldsymbol{3}}}$ |
|  | $p_{i}^{A_{3}A_{3}}$ | ***two copies*** of $A_{3}$  *(ie.* $A_{3}A_{3})$ |  |

If one of the alternate alleles is the major allele, then updating the allele dosage such that the baseline allele matches with the other allele is required to account for all the imputation probabilities at all the biallelic SNP variables for that multi-allelic SNP. Below, we describe the pseudo-algorithm implemented for multi-allelic SNPs to check if the major allele is the baseline allele and calculate the allele dosage for the new alternate allele if the baseline does not match.

**Pseudo-algorithm used in *recodeVCF* to recode multi-allelic SNPs**

**Input files:** **unfiltered** VCF file (type 4.0) including all the imputation probabilities for all SNPs in the region tested (or chromosome)

For each multi-allelic SNP with *M* alleles, including the baseline allele (can be for example the major allele or reference allele from the reference sequence genome) and *M-1* alternate alleles. The illustrative example from **Table S7**, has *M*=3 alleles.

**For *each* multi-allelic SNP,**

1) Compute the allele frequency of the **baseline allele**, noted $A_{M}$, with:

f($A_{M})=1-\sum_{m=2}^{M} f(A_{m})$

2) rank the allele by frequencies:

$\boldsymbol{f}\left( \boldsymbol{A}_{\boldsymbol{1}}^{\boldsymbol{*}} \right)>$…> $f\left( A_{m}^{*} \right)>\ldots>f\left( A_{M}^{*} \right)$

Where $A_{m}^{*}$ with 1 $\leq m\leq M$ denote the alleles ranked by frequency

(ie. $A_{1}^{*}$ correspond to the *most* frequent allele, $A_{M}^{*}$ to the *least* frequent)

3) **If** { $A_{1}^{*}$= $A_{M}$},

**then** { no update of the allele dosage ; **go** to the next multiallelic SNP }

**else {**

New BASELINE = $\boldsymbol{A}_{\boldsymbol{1}}^{\boldsymbol{*}}$

New ALT = $A_{M}$ & calculate the dosage of “New ALT” for each individual *i:*

$$d_{i} \left( A_{M} \right)=2(1-\sum_{m=2}^{M} \sum_{\begin{aligned} m^{'}=2 \\ m'\neq m \end{aligned}}^{M} p_{i}^{A_{m}A_{m}'})-\sum_{m=1}^{M} p_{i}^{A_{m}A_{m}}$$

Where:

- $p_{i}^{A_{m}A_{m}'}$ are the imputation probabilities for individual *i*, for all possible heterozygous genotypes for allele A*_m_* at the multi-allelic SNP considered
- $p_{i}^{A_{m}A_{m}}$ are the imputation probabilities for individual *i*, for all the homozygous genotypes A_m_A_m_ at the multi-allelic SNP considered

**go** to the next multi-allelic SNP

For the multi-allelic SNPs where the allele frequency and dosage values have been updated (and all other SNPs), we also report the machr2 imputation quality score calculated as the ratio of the empirically observed variance of the allele dosage to the expected binomial variance at Hardy-Weinberg equilibrium^24^, for each SNP (with MAF, noted *p*) as:

$Mach r^{2}=\left\{ \begin{aligned} \frac{\frac{\sum_{i=1}^{N} {d_{i}}^{2}}{N}-\left( \frac{\sum_{i=1}^{N} d_{i}}{N} \right)^{2}}{2p (1-p)}\mathrm{when}p \epsilon]0,1[ \\ 1 \mathrm{when}p=0 or p=1 \end{aligned} \right.$

This quality score can be used to filter the multiallelic SNPs that have been recoded, as well as all other biallelic SNPs.

##### 2.2.3 Description of outputs

- *Data* output, with dosage for bi-allelic & multi-allelic SNPs in region VCF input file, where multi-allelic SNPs are recoded such that their baseline allele is the major allele (and covariates if applicable)
- *SNPinfo* output including columns "chr", "pos", "variant","ref", "alt", "region", "start.bp", "end.bp", "multialSNP": binary variable=0, if the SNP is bi-allelic (two alleles), or =1 otherwise (if more than two alleles), "multialSNP.rec": indicates if the multi-allelic SNP has been recoded (such that baseline allele = major allele), "freq.alt": frequency of the alternate allele (minor allele), “machr_r2”: mach imputation quality score calculated for all the SNPs (can be used to filter multiallelic SNPs and all other SNPs that have been recoded such that their major allele is the baseline allele).

### Supplementary Information 3. Usage Case: Application of RegionScan in the DCCT/EDIC genetic study

#### 3.1 DCCT/EDIC analysis methods

To illustrate RegionScan capabilities, we conducted genome-wide region-level association analysis for low-density lipoprotein-cholesterol (LDL-C), measured at baseline in *N*=1340 individuals of European-ancestry from the DCCT/EDIC Genetic study^25–27^. Genome-wide genotyping was performed subsequently using Illumina 1M and HumanCoreExome Bead Arrays (Illumina, San Diego, CA, USA) and standard quality controls procedures were applied to individuals and genetic variants^27,28^. In this study, we use 1340 individuals genotyped with the HumanCoreExome Bead Array and ungenotyped autosomal SNPs imputed using 1000 Genomes^29^ data phase 3 (v5) and minimac3^30^ (v.1.0.13), as previously described^28^. Genomic positions are on build GRCh37.

To define regions for comprehensive analysis of the genome (including intergenic regions), we applied the BigLD^31^ algorithm (with default parameter values) implemented in R package gpart^32^ to each chromosome separately. In total, BigLD partitioned 5,687,207 bi-allelic autosomal SNPs (MAF > 0.05 and Mach imputation quality score R^2^ > 0.5) into 89,003 quasi-independent regions of varying sizes, adaptive to the chromosomal LD structure. We conducted genome-wide region-level analysis of these 89,001 regions based on a total of 5,682,968 biallelic SNP variables (ie 5,673,694 biallelic SNPs plus 9274 biallelic SNP variables corresponding to 4637 triallelic SNPs processed with recodeVCF) with LDL-c (untransformed) using the default region-level tests implemented in *regscan*, adjusted for sex, age, and sex by age interaction as covariates. Similarly to previous GWAS conducted in DCCT/EDIC individuals of European ancestry, we did not adjust for any ancestry PCs in this analysis. The main features of the DCCT/EDIC dataset analysis and options specified in *regscan* are summarized in **Table S9.** In **Table S10**, we report the number of regions and SNPs analyzed for each chromosome.

A total of 89,001 were analyzed with *regscan* (two regions encountered convergence issues in multiple regression models in chr16 and chr6, likely due to complex LD structures; requiring rerunning RegionScan for these two regions with a more stringent LD pruning). We identify regions that reach the genome-wide significant Bonferroni-corrected threshold of 5.62E-7 (which assumes independence between the 89,001 LD regions) for at least one of the region-level tests, and at a suggestive level of 1E-05. The Bonferroni-corrected region-level significance threshold was found to agree with an empirical permutation significance level in European-ancestry population of the Canadian Longitudinal Study on Aging (CLSA) genome-wide data^33^.

**Table S9.** Summary of DCCT/EDIC Genetic analysis

| # of individuals | 1340 |
| --- | --- |
| # of BigLD regions analyzed | 89,001 |
| # of variants in regions | 5,682,968 HumanCoreExome genotyped  & 1000G-imputed bi-allelic & tri-allelic SNP variables |
| Trait | LDL-C (at baseline) |
| Covariates | Sex, age, sex by age interaction |

**Table S10.** Number of regions identified by BigLD in DCCT/EDIC analyzed with *regscan* and number of variants in regions out of the 5,682,968 SNP variables

| **chr** | **# regions** | **Region size in bp**  **Mean [Min ; Max]** | **#SNPs within regions^1^** | **#SNPs analyzed^1,2^** |
| --- | --- | --- | --- | --- |
| 1 | 7,085 | 28,910 [2 ; 1,159,157] | 434,442 | 130,554 |
| 2 | 7,075 | 31,330 [2 ; 808,451] | 477,995 | 152,044 |
| 3 | 5,899 | 31,172 [2 ;1314,045] | 408,142 | 126,468 |
| 4 | 5,768 | 30,639 [2 ; 703,033] | 420,633 | 118,029 |
| 5 | 5,300 | 31,523 [2 ; 1,022,494] | 364,563 | 110,959 |
| 6 | 5,220 | 30,320 [2 ; 711,485] | 387,425 | 111,695 |
| 7 | 4,996 | 28,743 [2 ; 1,309,785] | 336,169 | 105,481 |
| 8 | 4,653 | 28,676 [2 ; 1,522,918] | 318,786 | 102,738 |
| 9 | 4,074 | 24,949 [2 ; 631,503] | 244,427 | 87,388 |
| 10 | 4,465 | 27,079 [2 ; 902,714] | 294,838 | 92,182 |
| 11 | 4,236 | 29,133 [2 ; 1,2345,539] | 289,324 | 87,981 |
| 12 | 4,252 | 28,834 [2 ; 1,013,631] | 276,420 | 86,484 |
| 13 | 3,232 | 27,981 [2 ; 689,896] | 212,951 | 64,863 |
| 14 | 2,898 | 27,896 [2 ; 596,532] | 183,245 | 58,673 |
| 15 | 2,886 | 24,630 [2 ; 761,812] | 158,021 | 55,364 |
| 16 | 3,332 | 20,187 [2 ; 498,792] | 170,822 | 66,849 |
| 17 | 2,919 | 23,684 [2 ; 777,204] | 146,123 | 49,213 |
| 18 | 2,830 | 24,518 [2 ; 542,101] | 160,707 | 52,487 |
| 19 | 2,548 | 19,289 [2 ; 587,416] | 122,236 | 41,450 |
| 20 | 2,474 | 22,075 [2 ; 362,193] | 124,006 | 43,439 |
| 21 | 1,330 | 22,844 [2 ; 362,193] | 79,143 | 25,832 |
| 22 | 1,531 | 19,173 [2 ; 509,158] | 72,550 | 26,412 |
| **Total** | **89,003^1^** |  | **5,682,968** | **1,796,585** |

^1^Based on 89,001 regions analyzed with *regscan* (analysis of two regions failed because of convergence issues on chr16 and chr6).

^2^after default pruning on MAF>0.05 and LD (SNP correlation > 0.99) within regions by *regscan*.

**Fig S3.** Distribution of region sizes (# of SNPs analyzed per region) among the 89,001 regions analyzed genome-wide

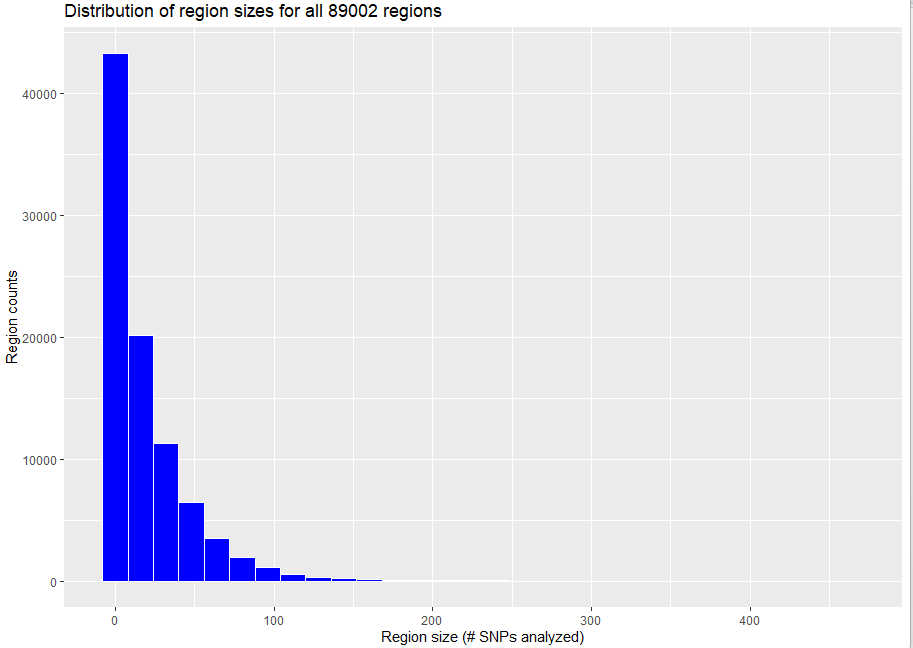

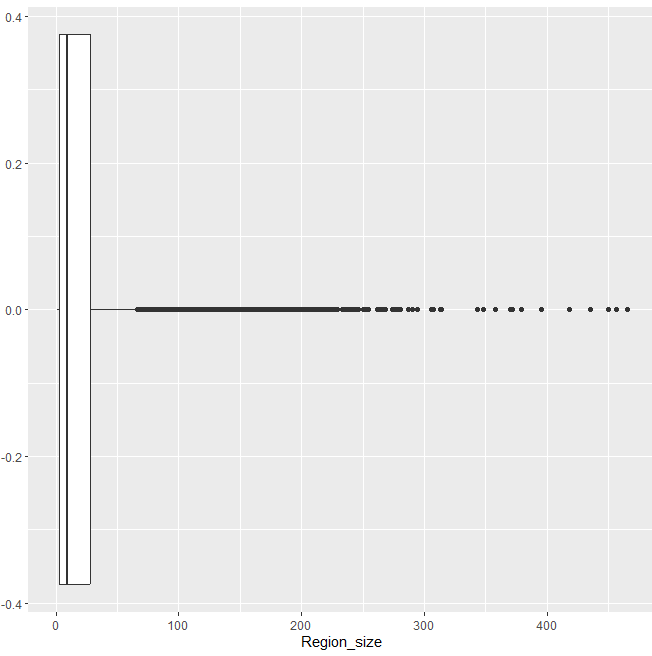

#### 3.2 RegionScan results

In **Fig S4**, we illustrate the genome-wide region-level results for the example of the MLC test, in comparison to the single-SNP results. Out of the 89,001 regions analyzed, we detect 5 regions around the *APOE* gene that meet the genome-wide region level significance threshold of 5.62E-7 (and 17 single-SNP tests with P≤5E-8), and two suggestive signals around the *LDLR* gene with P≤ 1E-5 ( 42 single-SNP tests with P≤1E-5, none of them met the genome-wide significance level); these two loci were reported by previous large scale meta-analyses (ref). Sections 3.2.2 and 3.2.3, subsequently illustrate the results for extended loci around the detected regions in *APOE* locus (chr19: 45257201- 45436657) and *LDLR* (chr19: 11071560- 11229577) loci. Outputs produced by *regscan* for the regions in these two loci are provided in Supplementary Tables.

**Fig S4.** Example of “Miami plot” produced by the utility function *MiamiPlot* based on the region-level MLC test P-values (top panel) and variant-level (bottom panel) P-values (-log 10). On the top panel, we show the region-level results for 89,001 regions analyzed, and **1,796,585** SNPs analyzed (after pruning) in these regions. The signals highlighted in orange meet the genome-wide significance criteria: of 5.6E-07 for region-level tests (dashed line on the top plot), of 5E-08 for single-SNP tests (dashed line on the bottom plot).

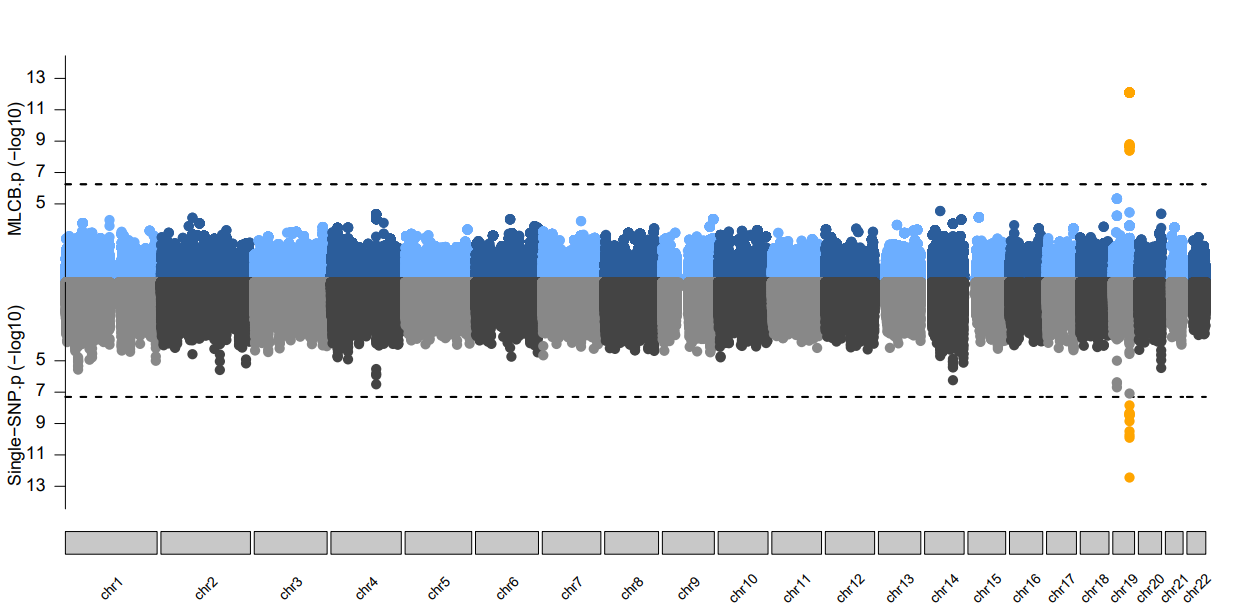

**Table S11.** Genomic control values for all 89,001 regions analyzed, reported by the *QQplot* function for three example of region-level tests

| **Region-level tests** | **Genomic control** |
| --- | --- |
| Wald | 1.026 |
| MLC | 1.002 |
| PC80 | 1.00 |
| SKATO | 0.955 |

**Fig S5.** QQ-plots for all 89,001 regions analyzed from the four region-level tests from **Table S11**

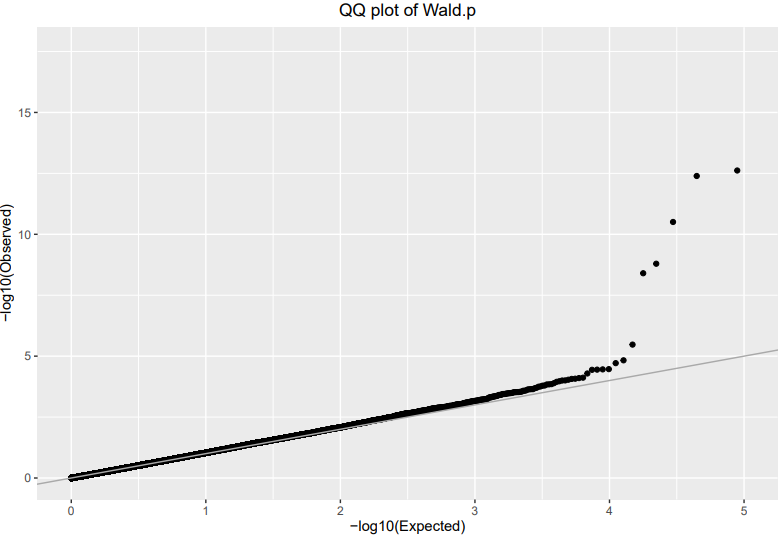

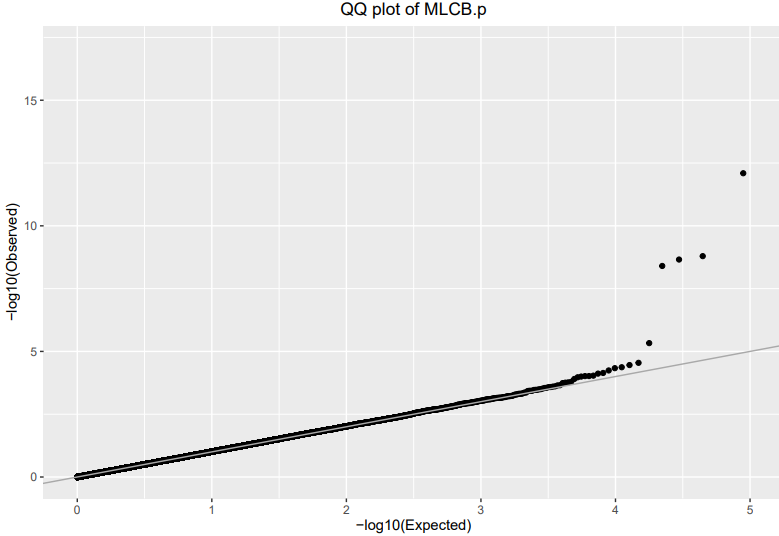

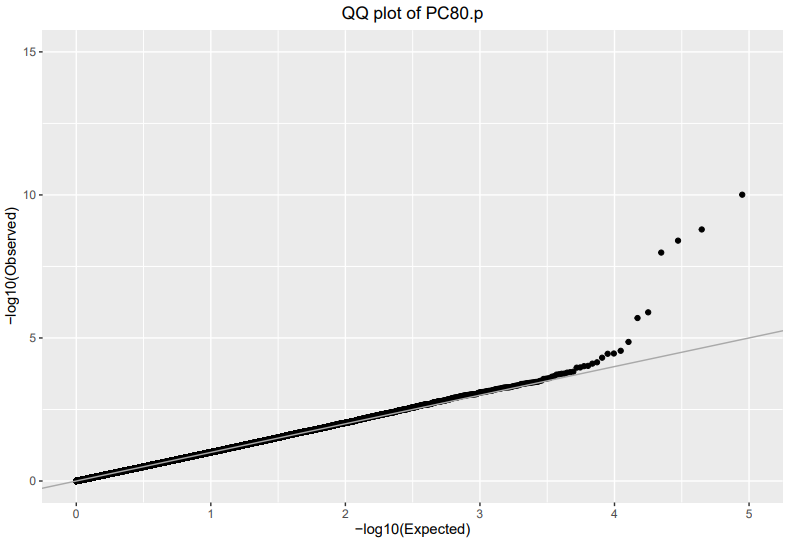

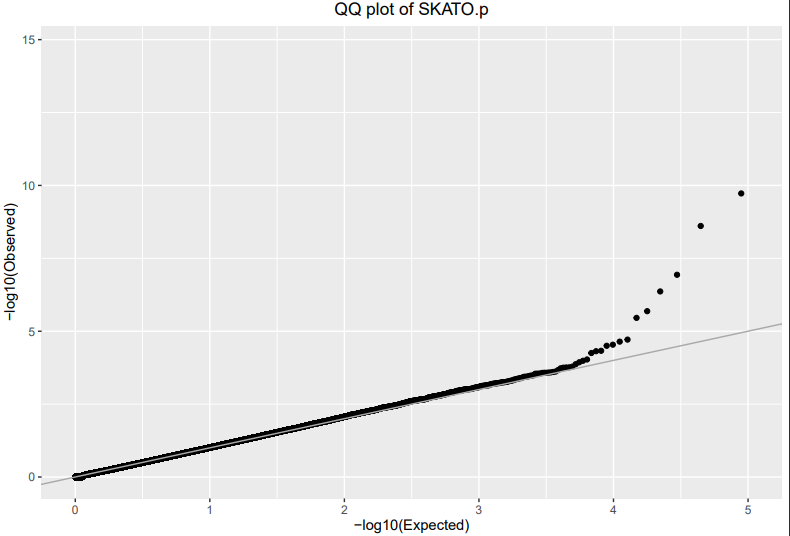

**Table S12.** Regions exhibiting association signals on chr19 (+ neighboring regions) extracted from the region-level output.

| **chr** | **region** | **start.bp** | **end.bp** | **nSNPs** | **nSNPs.kept** | **maxVIF** | **Wald** | | **MLC** | | **PC80** | | **SKATO.p** | **uMinP.p** |
| --- | --- | --- | --- | --- | --- | --- | --- | --- | --- | --- | --- | --- | --- | --- |
|  |  |  |  |  |  |  | df | P | df | P | df | P |  |  |
| 19 | 619 | 10961273 | 10962440 | 3 | 2 | 35.1 | 2 | 2.6E-01 | 1 | 1.9E-01 | 1 | 1.9E-01 | 1.9E-01 | 1.7E-01 |
| 19 | 620 | 10962974 | 11069264 | 120 | 40 | 1480.5 | 40 | 9.8E-02 | 6 | 9.2E-02 | 3 | 2.4E-01 | 2.8E-01 | 1.2E-02 |
| 19 | 621 | 11069908 | 11071395 | 6 | 2 | 1.2 | 2 | 2.4E-01 | 2 | 2.4E-01 | 2 | 2.4E-01 | 1.8E-01 | 9.2E-02 |
| 19 | 622 | 11071560 | 11157423 | 106 | 30 | 16605.8 | 30 | 8.9E-02 | 3 | 1.5E-01 | 2 | 1.7E-01 | 1.2E-01 | 2.3E-03 |
| 19 | 623 | 11159076 | 11185014 | 67 | 23 | 772.9 | 23 | 6.2E-03 | 7 | 4.1E-03 | 4 | 1.1E-01 | 7.6E-02 | 1.2E-02 |
| 19 | 624 | 11185919 | 11202306 | 51 | 8 | 86.8 | 8 | 1.3E-04 | 3 | **4.4E-06** | 2 | **1.2E-06** | **3.3E-06** | **1.9E-07** |
| 19 | 625 | 11205975 | 11214533 | 20 | 7 | 105.1 | 7 | 1.3E-03 | 4 | 1.3E-04 | 2 | 2.0E-05 | 2.8E-04 | 1.1E-05 |
| 19 | 626 | 11216561 | 11228745 | 26 | 14 | 755.1 | 14 | 2.8E-01 | 4 | 8.9E-02 | 2 | 1.7E-02 | 1.1E-02 | 2.1E-03 |
| 19 | 627 | 11228783 | 11229577 | 5 | 2 | 1.2 | 2 | 6.9E-03 | 2 | 6.9E-03 | 2 | 6.9E-03 | 9.5E-03 | 3.5E-03 |
| 19 | 628 | 11229765 | 11250396 | 105 | 35 | 20886.1 | 35 | 1.2E-02 | 8 | 1.5E-01 | 3 | 1.1E-02 | 4.2E-03 | 9.7E-04 |
| 19 | 629 | 11253310 | 11262319 | 25 | 21 | 176.5 | 21 | 8.7E-02 | 9 | 6.7E-02 | 4 | 1.2E-02 | 1.4E-01 | 2.1E-03 |
| 19 | 630 | 11262477 | 11280183 | 72 | 26 | 1681.6 | 26 | 3.4E-01 | 6 | 1.5E-01 | 3 | 4.9E-02 | 4.3E-02 | 4.2E-03 |
| 19 | 1683 | 45257201 | 45320386 | 60 | 31 | 347.27 | 31 | 1.6E-01 | 7 | 2.2E-01 | 4 | 1.1E-01 | 5.2E-01 | 1.1E-01 |
| 19 | 1684 | 45322744 | 45353261 | 93 | 49 | 4365.55 | 49 | 1.5E-02 | 9 | 3.7E-03 | 4 | 6.7E-03 | 1.3E-02 | 4.1E-05 |
| 19 | 1685 | 45354044 | 45354296 | 3 | 2 | 1.28 | 2 | 3.6E-05 | 2 | 3.6E-05 | 2 | 3.6E-05 | 2.9E-04 | 2.9E-05 |
| 19 | 1686 | 45355595 | 45359706 | 19 | 9 | 528.08 | 9 | 7.6E-03 | 4 | 5.8E-03 | 3 | 3.5E-02 | 1.0E-02 | 8.0E-04 |
| 19 | 1687 | 45360573 | 45360762 | 3 | 3 | 5.51 | 3 | 1.8E-01 | 2 | 3.1E-01 | 2 | 3.2E-01 | 5.2E-01 | 1.3E-01 |
| 19 | 1688 | 45360967 | 45360968 | 2 | 1 | 1.00 | 1 | 6.3E-01 | 1 | 6.3E-01 | 1 | 6.3E-01 | 6.3E-01 | 6.3E-01 |
| 19 | 1689 | 45361224 | 45384116 | 98 | 40 | 699.20 | 40 | 2.2E-04 | 9 | 7.0E-05 | 4 | 8.9E-01 | 3.5E-01 | 6.8E-03 |
| 19 | 1690 | 45385759 | 45415935 | 61 | 38 | 356.11 | 38 | **5.6E-14** | 11 | **1.7E-12** | 5 | **1.4E-10** | **4.0E-06** | **6.1E-13** |
| 19 | 1691 | 45416178 | 45418790 | 6 | 4 | 58.30 | 4 | **2.1E-13** | 2 | 5.1E-02 | 2 | 1.0E-01 | **2.2E-07** | **1.8E-08** |
| 19 | 1692 | 45421254 | 45424514 | 9 | 6 | 47.82 | 6 | **2.6E-11** | 2 | **2.5E-09** | 2 | **1.2E-08** | **2.2E-10** | **1.7E-10** |
| 19 | 1693 | 45425178 | 45426792 | 3 | 2 | 1.04 | 2 | **1.9E-09** | 2 | **1.9E-09** | 2 | **1.9E-09** | **3.0E-09** | **9.6E-08** |
| 19 | 1694 | 45427125 | 45428234 | 3 | 2 | 1.12 | 2 | **5.3E-09** | 2 | **5.3E-09** | 2 | **5.3E-09** | **2.1E-08** | **4.6E-09** |
| 19 | 1695 | 45428459 | 45430280 | 3 | 3 | 4.68 | 3 | 4.0E-01 | 1 | 9.6E-01 | 1 | 9.8E-01 | 6.3E-01 | 5.1E-01 |
| 19 | 1696 | 45431453 | 45436657 | 11 | 4 | 53.66 | 4 | 1.1E-02 | 2 | 1.0E-02 | 2 | 8.9E-03 | 3.8E-03 | 6.4E-04 |

#### 3.3 Focus in two regions exhibiting association signals

##### 3.3.1 “APOE locus” (chr19: 45257201- 45436657)

**Fig S6.** LD (r2) heatmap of the 14 consecutive BigLD regions in chr19: 45257201- 45436657 (regions #1683 to #1696) produced using the LDheatmap function from the gpart package (for all biallelic SNPs satisfying QC and analyzed in BigLD). Protein coding gene positions (from GENCODE) are shown in Build 37.

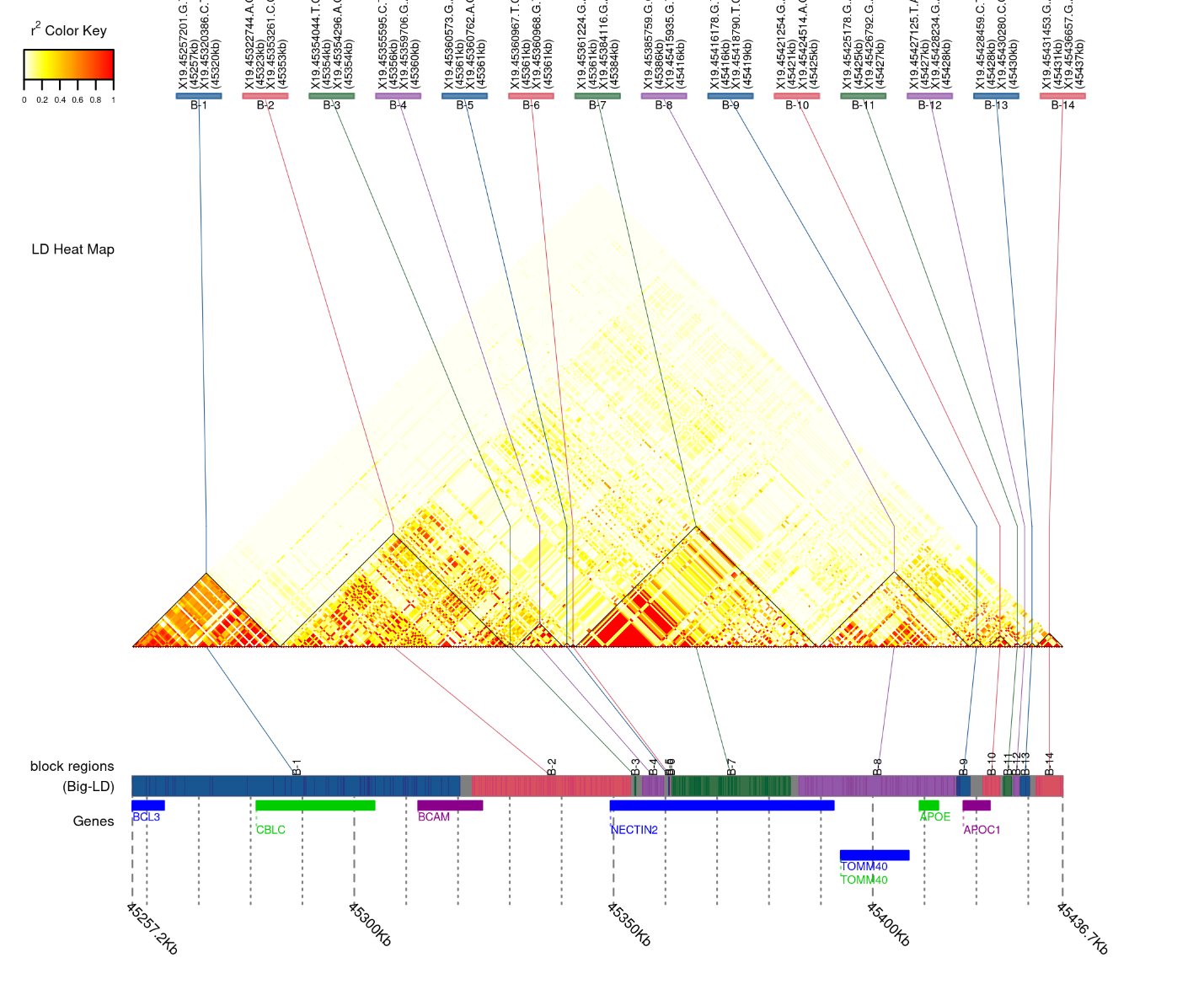

**Fig S7.** Multi-region association plot for regions # 1683 to 1696 in chr19. The *Y* axis shows -log10(*P*-values) for five region-level tests and all single-SNP tests. Dashed horizontal lines indicate GW significance thresholds: 5E-8 for single-SNP analysis (blue dotted line) and the genome-wide region-level significance threshold of 5.62E-7 (red dotted line). Region numbers appear at the bottom of the plots. In Panel (A.) the *X* axis shows the physical positions of the regions/SNPs, and region boundaries are indicated by shading in alternating regions. In Panel (B.) regions are represented with a fixed size to facilitate comparisons across regions; region test labels indicate the number of SNPs analyzed in each region.

(A).

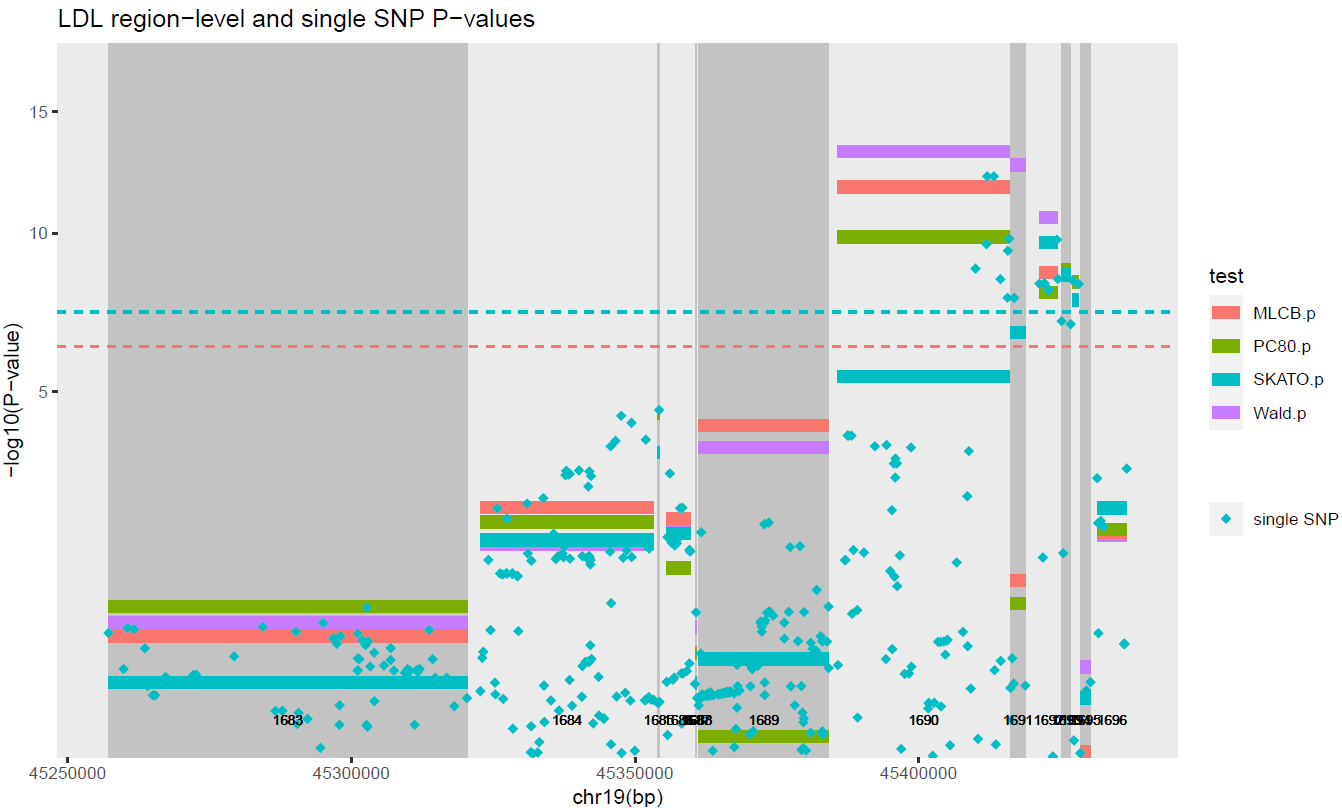

(B.)

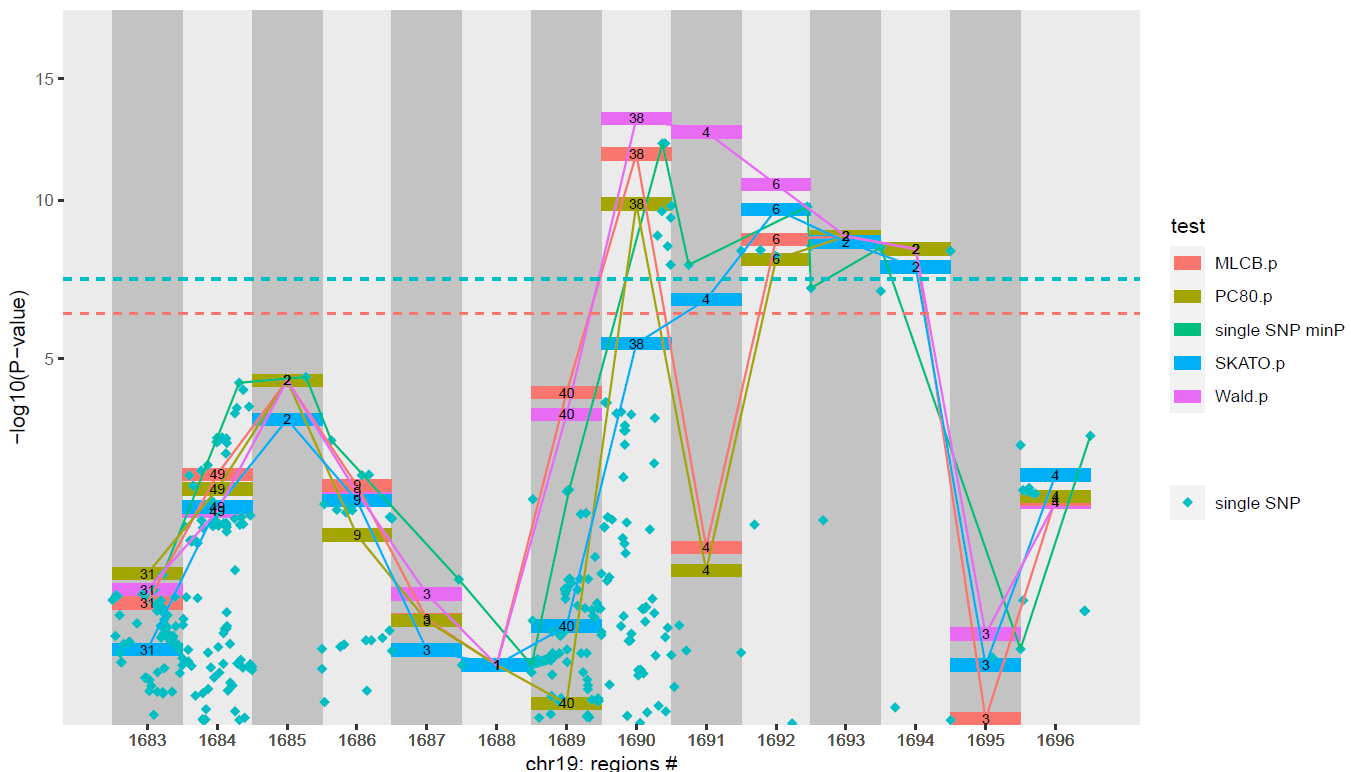

**Example: Detailed results for region #1690**

The region #1690 is identified as the top region genome-wide by at least three of the region-level tests (*P*<5.62E-7) and by single-SNP analysis (P<5E-8). This region overlaps *APOE* & *TOMM40* and includes two well-known *APOE* SNPs, rs429358 and rs7412, reported associated with LDL-C by previous genetic association studies^34,35^. The Wald region-level test has a smaller *P-*value than the three other region-level tests (**Table S13**). Bin-level statistics in region #1690 show no signal for LD bin2 (which includes rs429358), and a strong signal for LD bin3 (which includes rs7412). These results might result from the genetic architecture in this region and would require more investigations, which is beyond the purpose of this analysis to illustrate the application of the proposed *RegionScan* R package.

**Table S13.** Region-level test results in region #1690 (extract from the “region-level” output)

| **chr** | **region** | **start.bp** | **end.bp** | **nSNPs** | **nSNPs.kept** | **maxVIF** | **Wald** | | **MLC** | | **PC80** | | **SKATO.p** | **ACAT.p^1^** |
| --- | --- | --- | --- | --- | --- | --- | --- | --- | --- | --- | --- | --- | --- | --- |
|  |  |  |  |  |  |  | df | P | df | P | df | P |  |  |
| 19 | 1690 | 45385759 | 45415935 | 61 | 38 | 356.11 | 38 | **5.6E-14** | 11 | **1.7E-12** | 5 | **1.4E-10** | **4.0E-06** | **2.2E-13** |

^1^aggregated Cauchy association (ACAT) test^36^ calculated from the Wald, MLC and PC80 region-level test *P*-values.

**Table S14**. Bin-level test results in region #1690 (extract from the “bin-level” output)

| chr | region | bin | binstart.bfP.(bp) | binend.bfP.(bp) | binstart.afP.(bp) | binend.afP.(bp) | binsize.bfP | binsize.afP | deltaB | deltaB.se | deltaB.pvalue |
| --- | --- | --- | --- | --- | --- | --- | --- | --- | --- | --- | --- |
| 19 | 1690 | 1 | 45,385,759 | 45,410,444 | 45,385,759 | 45,410,444 | 22 | 12 | -0.52 | 0.34 | 1.2E-01 |
| 19 | 1690 | 2* | 45,387,459 | 45,415,935 | 45,387,459 | 45,415,935 | 19 | 10 | -0.26 | 0.34 | 4.4E-01 |
| 19 | 1690 | 3* | 45,412,079 | 45,415,640 | 45,413,233 | 45,415,640 | 4 | 2 | -11.93 | 1.99 | 2.1E-09 |
| 19 | 1690 | 4 | 45,387,034 | 45,389,174 | 45,387,057 | 45,389,174 | 4 | 3 | 2.69 | 1.18 | 2.2E-02 |
| 19 | 1690 | 5 | 45,398,633 | 45,408,564 | 45,398,633 | 45,408,564 | 3 | 3 | 0.41 | 0.94 | 6.6E-01 |
| 19 | 1690 | 6 | 45,402,718 | 45,403,924 | 45,402,718 | 45,402,718 | 2 | 1 | 2.87 | 3.33 | 3.9E-01 |
| 19 | 1690 | 7 | 45,394,211 | 45,413,366 | 45,394,211 | 45,413,366 | 2 | 2 | 0.95 | 1.29 | 4.6E-01 |
| 19 | 1690 | 8 | 45,404,691 | 45,414,451 | 45,404,691 | 45,414,451 | 2 | 2 | -4.19 | 2.01 | 3.7E-02 |
| 19 | 1690 | 9 | 45,389,224 | 45,389,224 | 45,389,224 | 45,389,224 | 1 | 1 | 2.08 | 2.92 | 4.8E-01 |
| 19 | 1690 | 10 | 45,408,628 | 45,408,628 | 45,408,628 | 45,408,628 | 1 | 1 | -0.07 | 2.78 | 9.8E-01 |
| 19 | 1690 | 11 | 45,413,576 | 45,413,576 | 45,413,576 | 45,413,576 | 1 | 1 | 5.33 | 3.10 | 8.5E-02 |

*bin2 includes rs429358 and bin3 rs7412; combination of alleles C at rs429358 and allele C at rs7412 is known as the APOE-e4 allele which has been reported associated with LDL-C and increased risk for heart disease.

**Table S15**. Single variant-level test results in region #1690 (extract from the “variant-level” output) for the 38 variants kept after LD pruning.

| chr | region | bin | variant | rsID | multiallelic | ref | alt | maf | MLC.  codechange | Single-SNP analysis | |
| --- | --- | --- | --- | --- | --- | --- | --- | --- | --- | --- | --- |
|  |  |  |  |  |  |  |  |  |  | sglm.  Beta | sglm.  Pvalue |
| 19 | 1690 | 1 | 19.45385759.G.C | rs3745150 | 0 | G | C | 0.41 | 0 | 1.02 | 3.9E-01 |
| 19 | 1690 | 1 | 19.45395266.G.A | rs157580 | 0 | G | A | 0.38 | 1 | -3.23 | 4.1E-03 |
| 19 | 1690 | 1 | 19.45395330.A.G | rs2075649 | 0 | A | G | 0.39 | 0 | 1.20 | 2.9E-01 |
| 19 | 1690 | 1 | 19.45396899.T.C | rs157584 | 0 | T | C | 0.49 | 0 | 0.07 | 9.5E-01 |
| 19 | 1690 | 1 | 19.45398716.A.C | rs157590 | 0 | A | C | 0.48 | 0 | -0.94 | 4.0E-01 |
| 19 | 1690 | 1 | 19.45401666.A.G | rs8106922 | 0 | A | G | 0.41 | 0 | -0.42 | 7.1E-01 |
| 19 | 1690 | 1 | 19.45401783.T.C | rs56290633 | 0 | T | C | 0.44 | 1 | -0.56 | 6.7E-01 |
| 19 | 1690 | 1 | 19.45405521.G.C | rs1305062 | 0 | G | C | 0.40 | 0 | -0.09 | 9.3E-01 |
| 19 | 1690 | 1 | 19.45407788.G.A | rs7259620 | 0 | G | A | 0.45 | 0 | -1.44 | 2.0E-01 |
| 19 | 1690 | 1 | 19.45408836.T.G | rs405509 | 0 | T | G | 0.48 | 1 | 4.08 | 2.7E-04 |
| 19 | 1690 | 1 | 19.45409167.C.G | rs440446 | 0 | C | G | 0.35 | 1 | -0.68 | 5.6E-01 |
| 19 | 1690 | 1 | 19.45410444.G.A | rs769450 | 0 | G | A | 0.40 | 0 | -0.14 | 9.0E-01 |
| 19 | 1690 | 2 | 19.45387459.C.G | rs12972156 | 0 | C | G | 0.14 | 0 | 6.15 | 1.2E-04 |
| 19 | 1690 | 2 | 19.45388500.A.G | rs283811 | 0 | A | G | 0.22 | 0 | 3.23 | 1.9E-02 |
| 19 | 1690 | 2 | 19.45392254.C.T | rs6857 | 0 | C | T | 0.17 | 0 | 5.45 | 2.1E-04 |
| 19 | 1690 | 2 | 19.45395844.G.A | rs34095326 | 0 | G | A | 0.11 | 0 | 5.76 | 9.8E-04 |
| 19 | 1690 | 2 | 19.45396144.C.T | rs11556505 | 0 | C | T | 0.14 | 0 | 5.41 | 5.0E-04 |
| 19 | 1690 | 2 | 19.45396219.C.T | rs157582 | 0 | C | T | 0.22 | 0 | 2.50 | 6.2E-02 |
| 19 | 1690 | 2 | 19.45396665.G.T | rs17855927 | 0 | G | T | 0.21 | 0 | 3.13 | 2.3E-02 |
| 19 | 1690 | **2** | 19.45410002.G.A | rs769449 | 0 | G | A | 0.12 | 0 | 10.03 | **1.8E-09** |
| 19 | 1690 | **2** | 19.45411941.T.C | rs429358 | 0 | T | C | 0.15 | 0 | 9.85 | **2.5E-10** |
| 19 | 1690 | **2** | 19.45415935.G.T | rs7256200 | 0 | G | T | 0.12 | 0 | 10.91 | **1.6E-10** |
| 19 | 1690 | **3** | 19.45413233.G.T | rs1065853 | 0 | G | T | 0.08 | 0 | -14.38 | **6.2E-13** |
| 19 | 1690 | **3** | 19.45415640.G.A | rs445925 | 0 | G | A | 0.11 | 0 | -9.84 | **1.7E-08** |
| 19 | 1690 | 4 | 19.45387057.A.G | rs283809 | 0 | A | G | 0.05 | 0 | -6.05 | 2.7E-02 |
| 19 | 1690 | 4 | 19.45388241.T.G | rs283810 | 0 | T | G | 0.08 | 0 | -3.39 | 1.3E-01 |
| 19 | 1690 | 4 | 19.45389174.T.A | rs283813 | 0 | T | A | 0.08 | 0 | -3.44 | 1.2E-01 |
| 19 | 1690 | 5 | 19.45398633.G.C | rs11668327 | 0 | G | C | 0.17 | 0 | -5.89 | 2.2E-04 |
| 19 | 1690 | 5 | 19.45406673.G.A | rs10119 | 0 | G | A | 0.29 | 0 | 2.83 | 2.9E-02 |
| 19 | 1690 | 5 | 19.45408564.A.T | rs449647 | 0 | A | T | 0.18 | 0 | -5.23 | 2.3E-03 |
| 19 | 1690 | 6 | 19.45402718.G.C | rs35568738 | 0 | G | C | 0.05 | 0 | -1.09 | 6.7E-01 |
| 19 | 1690 | 7 | 19.45394211.C.T | rs76692773 | 0 | C | T | 0.09 | 0 | 2.02 | 3.5E-01 |
| 19 | 1690 | 7 | 19.45413366.T.C | rs1081106 | 0 | T | C | 0.09 | 0 | 1.86 | 4.2E-01 |
| 19 | 1690 | 8 | 19.45404691.A.G | rs405697 | 0 | A | G | 0.26 | 0 | -1.31 | 3.2E-01 |
| 19 | 1690 | 8 | 19.45414451.T.C | rs439401 | 0 | T | C | 0.35 | 0 | -1.05 | 3.6E-01 |
| 19 | 1690 | 9 | 19.45389224.A.G | rs283814 | 0 | A | G | 0.06 | 0 | -0.76 | 7.8E-01 |
| 19 | 1690 | 10 | 19.45408628.T.C | rs769446 | 0 | T | C | 0.09 | 0 | -2.45 | 2.7E-01 |
| 19 | 1690 | 11 | 19.45413576.C.T | rs75627662 | 0 | C | T | 0.20 | 0 | 0.12 | 9.3E-01 |

rs1065853 is a proxy of rs7412; combination of alleles at rs429358 (C or G) and rs7412 (C or G) is known as the APOE-e4 allele which has been associated with LDL-C and increased risk for heart disease.

**Fig S8.** Visualization of the correlation structure **within region #1690** for the SNPs analyzed (after pruning & recoding), with SNPs ordered by physical position (A), and (B) SNPs ordered by LD bins (SNPs part of the same LD bin are indicated by the same color bands). Panel (C.) shows the physical position of the SNPs in the region #1690 by MLC LD bins (Y axis); the colors of the LD bins match those shown in Panel (B.), SNPs pruned out are shown in grey (with counts before/after pruning).

(A.)

(B.)

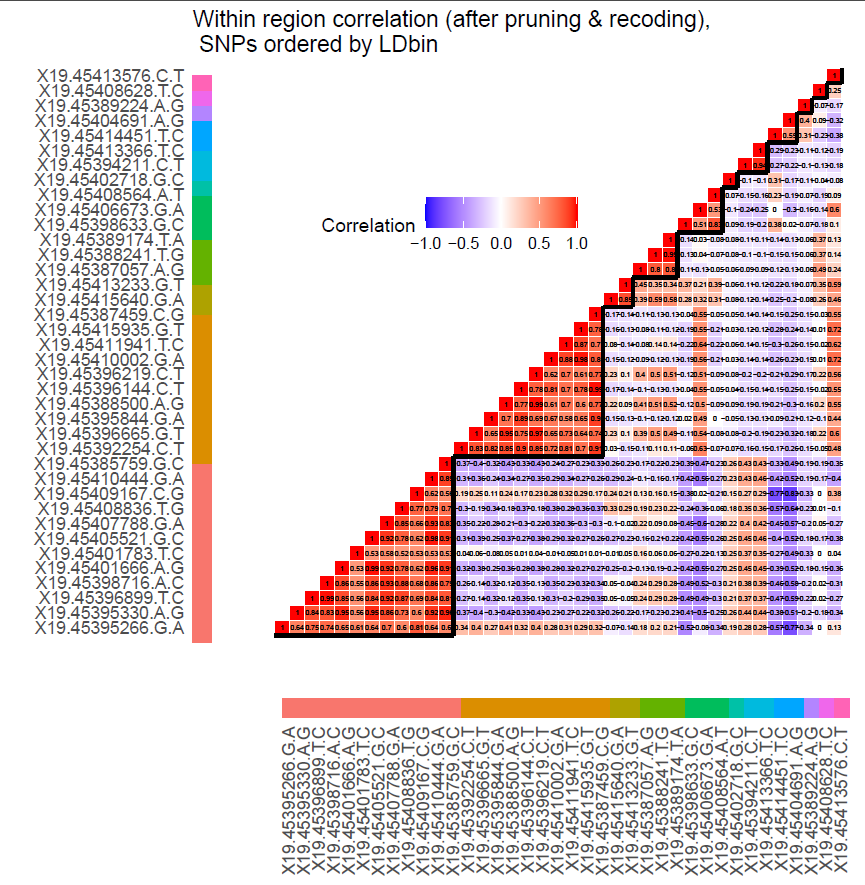

(C.)

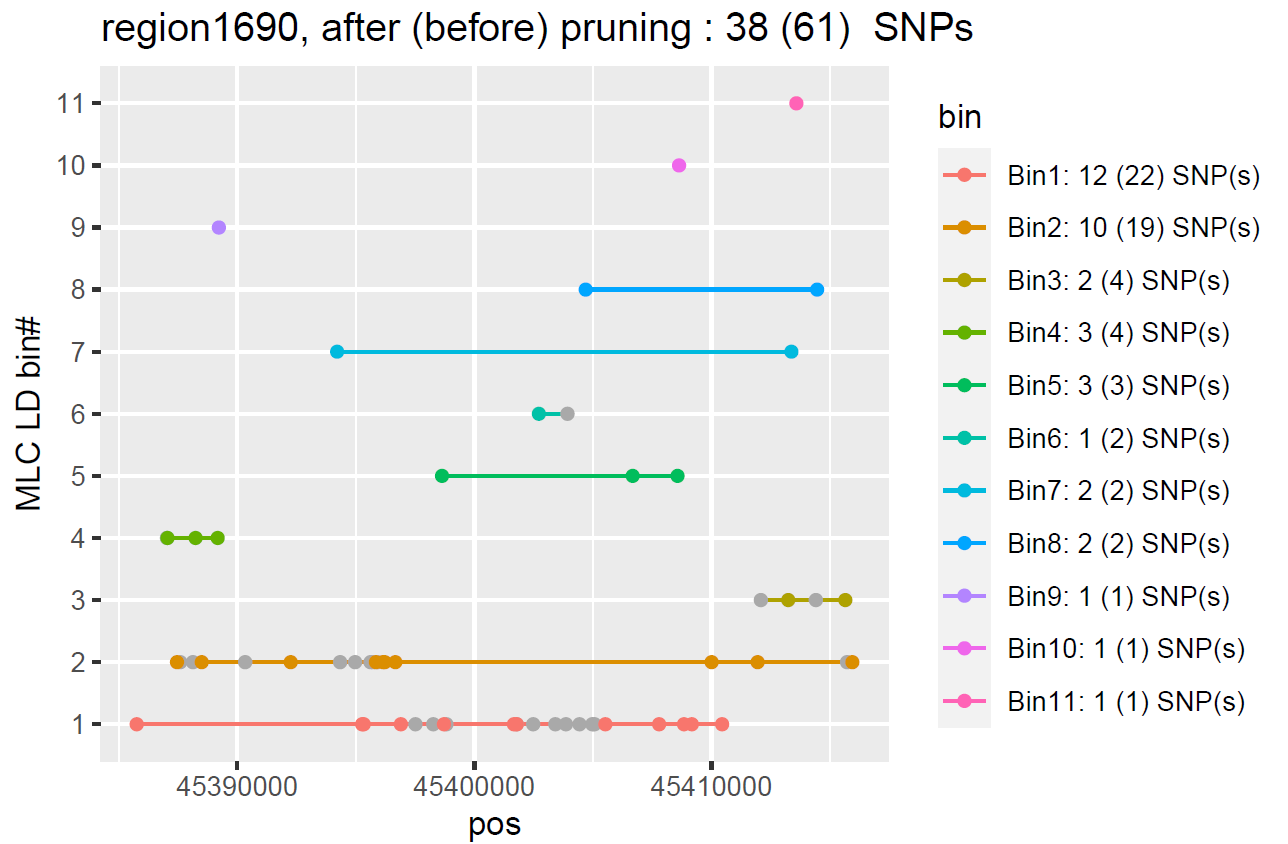

##### 3.3.2 “LDLR locus” (chr19: 11071560- 11229577)

**Figure S9.** Visualization of LD blocks within **chr19: 11071560- 11229577**; plots produced by LDheatmap function from gpart package (for all unfiltered biallelic SNPs used in BigLD)

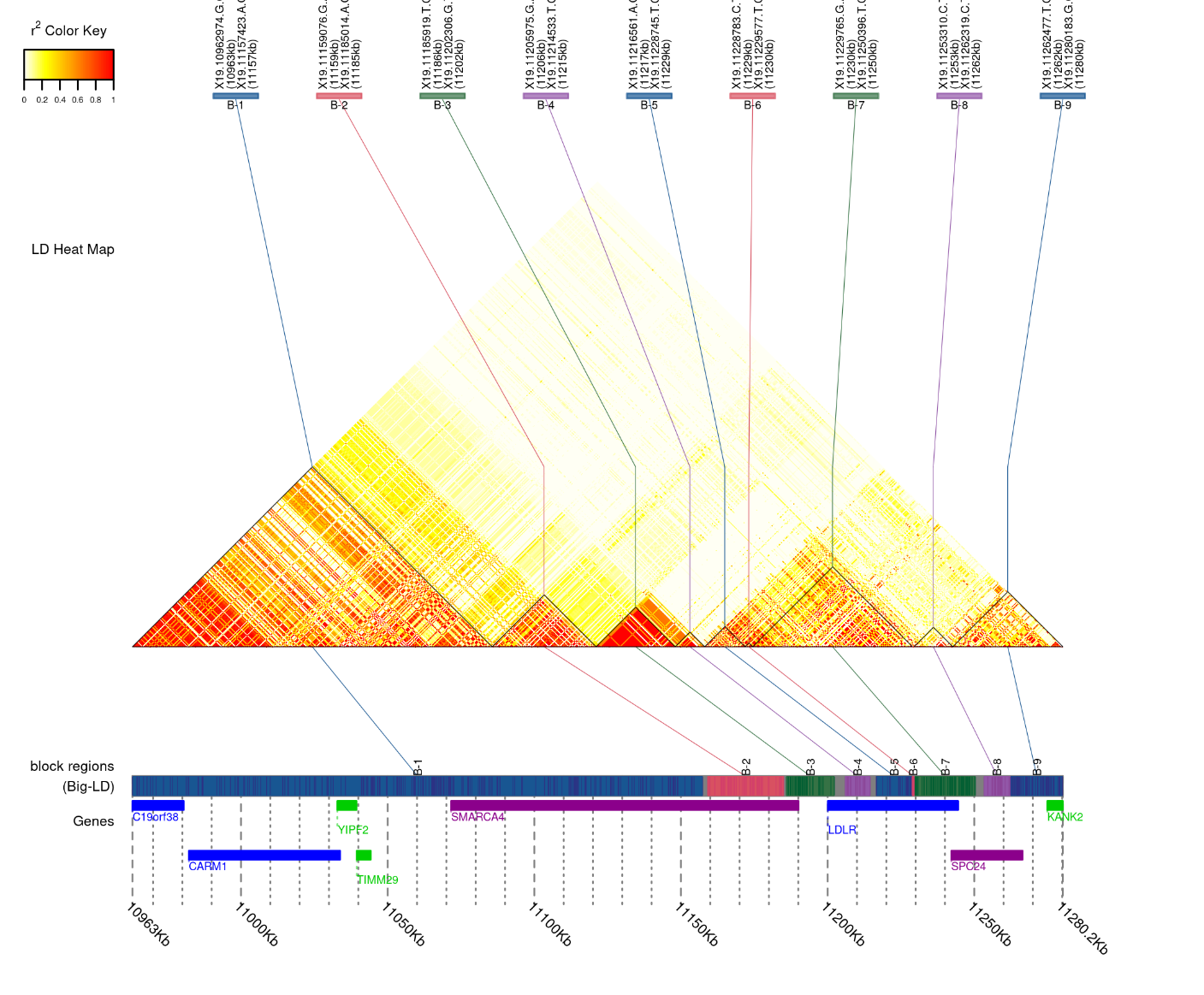

**Fig S10.** Multi-region association plot for regions #620 to 630 in chr19. The *Y* axis shows -log10(P-values) for five region-level tests and all single-SNP tests. Dashed horizontal lines indicate GW significance thresholds: 5E-8 for single-SNP analysis (blue dotted line) and the genome-wide region-level significance threshold of 5.62E-7 (red dotted line) for region-level. Region numbers appear at the bottom of the plots. In Panel (A.) the *X* axis shows the physical positions of the regions/SNPs, and region boundaries are indicated by shading in alternating regions. In Panel (B.) regions are represented with a fixed size to facilitate comparisons across regions; region test labels indicate the number of SNPs analyzed in each region.

(A).

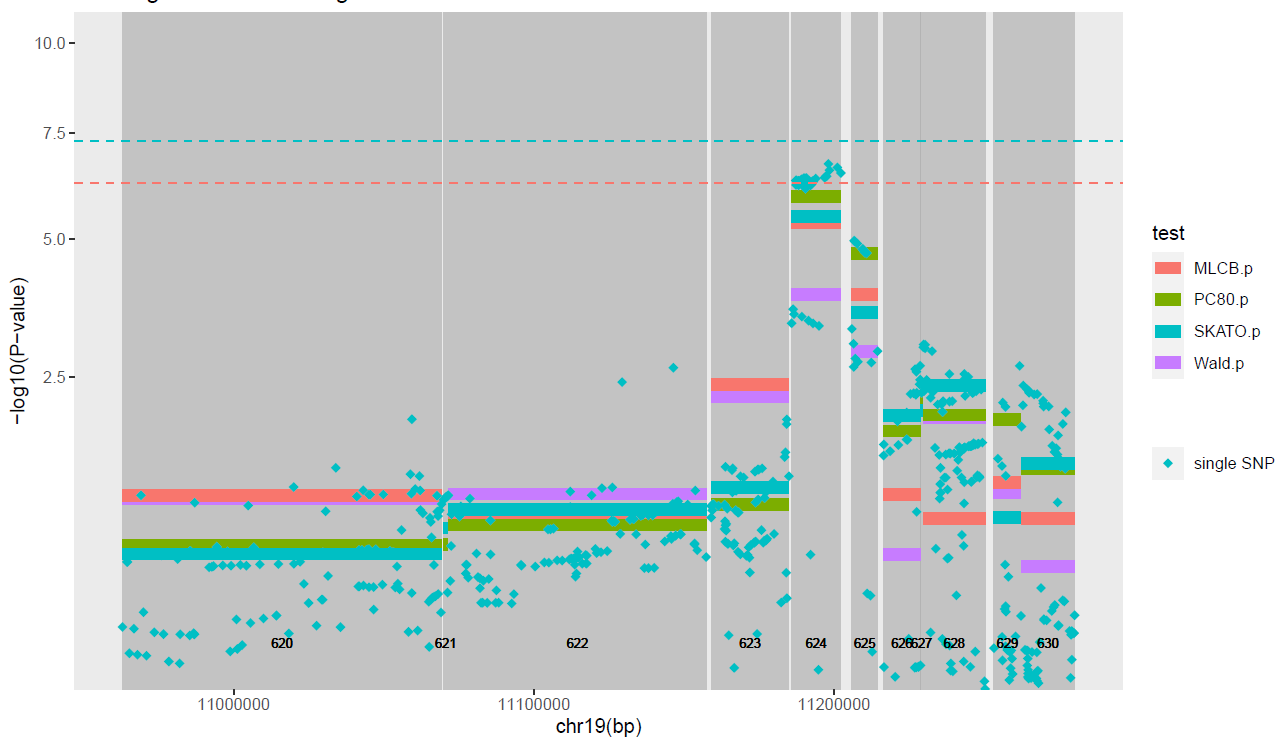

(B.)

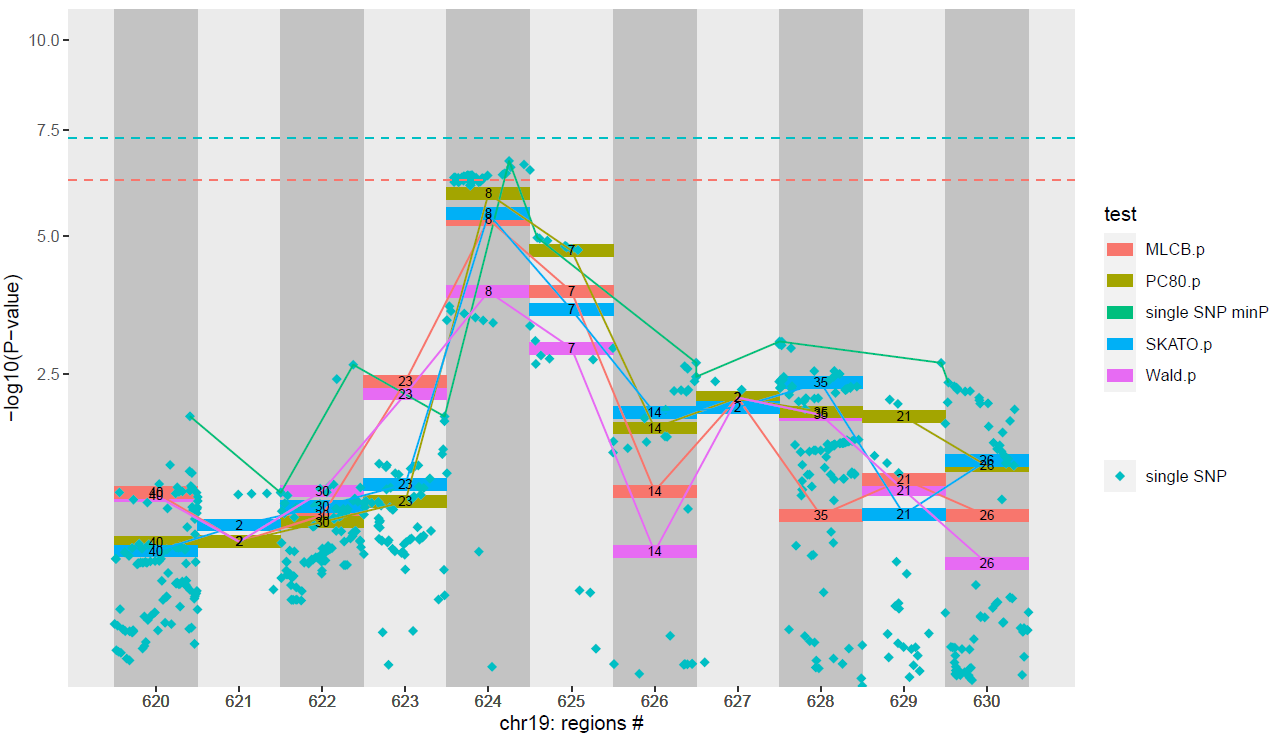

**Example: Detailed results for region #624**

Region #624 reached the genome-wide significance level of 5.62E-7 for region-level tests based on the uncorrected MinP test, and a suggestive level of *P*<5E-7 for reduced-df region-level tests (MLC, PC80), and variance-like component score test SKATO (**Table S16**). MLC, PC80 and SKATO exhibited stronger association with LDL-C than the Generalized Wald region-level test. Region #624 overlaps with rs6511720, reported consistently associated with LDL-C at the genome-wide significance level by previous GWAS^37–39^. Bin-level results showed a significant association for LD bin1 (**Table S17**) which includes 3 SNPs retained for the analysis with *P*<5E-7 in single-SNP analysis out of which rs6511720 (**Table S18**).

**Table S16.** Region-level test results for region #624

| **chr** | **region** | **start.bp** | **end.bp** | **nSNPs** | **nSNPs.**  **kept** | **maxVIF** | **Wald** | | **MLC** | | **PC80** | | **SKATO.p** | **ACAT.p** |
| --- | --- | --- | --- | --- | --- | --- | --- | --- | --- | --- | --- | --- | --- | --- |
|  |  |  |  |  |  |  | df | P | df | P | df | P |  |  |
| 19 | 624 | 11185919 | 11202306 | 51 | 8 | 86.8 | 8 | 1.3E-04 | 3 | 4.4E-06 | 2 | 1.2E-06 | 3.3E-06 | 2.9E-06 |

**Table S17.** Bin-level results in region #624

| **chr** | **region** | **bin** | **binstart.**  **bfP.bp** | **binend.**  **bfP.bp** | **binstart.**  **afP.bp** | **binend.**  **afP.bp** | **binsize.**  **bfP** | **binsize.**  **afP** | **NSNPs.**  **kept** | **deltaB** | **deltaB.se** | **deltaB.**  **pvalue** |
| --- | --- | --- | --- | --- | --- | --- | --- | --- | --- | --- | --- | --- |
| 19 | 624 | 1 | 11185919 | 11202306 | 11185919 | 11202306 | 49 | 6 | 6 | -1.59 | 0.31 | 2.8E-07 |
| 19 | 624 | 2 | 11192226 | 11192226 | 11192226 | 11192226 | 1 | 1 | 1 | 6.55 | 2.65 | 1.4E-02 |
| 19 | 624 | 3 | 11194823 | 11194823 | 11194823 | 11194823 | 1 | 1 | 1 | -1.27 | 3.30 | 7.0E-01 |

**Table S18.** Variant-level test results for all 8 variants analyzed region #624 (kept after LD pruning)

| **chr** | **region** | **bin** | **variant** | **rsID** | **multiallelic** | **ref** | **alt** | **maf** | **MLC.codechange** | **Single-SNP results** | |
| --- | --- | --- | --- | --- | --- | --- | --- | --- | --- | --- | --- |
|  |  |  |  |  |  |  |  |  |  | **sglm.**  **beta** | **sglm.**  **pvalue** |
| 19 | 624 | 1 | X19.11185919.T.C | rs10423733 | 0 | T | C | 0.17 | 0 | -5.5 | 4.3E-04 |
| 19 | 624 | 1 | X19.11186412.C.T | rs9305019 | 0 | C | T | 0.16 | 0 | -5.8 | 2.4E-04 |
| 19 | 624 | 1 | X19.11193091.T.G | rs112552009 | 0 | T | G | 0.10 | 0 | -9.2 | 5.2E-07 |
| 19 | 624 | 1 | X19.11195030.A.G | rs11668477 | 0 | A | G | 0.19 | 0 | -5.0 | 4.7E-04 |
| 19 | 624 | 1 | X19.11198187.C.T | rs17248720 | 0 | C | T | 0.11 | 0 | -9.6 | 1.9E-07 |
| **19** | **624** | **1** | **X19.11202306.G.T** | **rs6511720** | **0** | **G** | **T** | **0.10** | **0** | **-9.3** | **3.2E-07** |
| 19 | 624 | 2 | X19.11192226.T.C | rs7249753 | 0 | T | C | 0.06 | 0 | 2.7 | 2.8E-01 |
| 19 | 624 | 3 | X19.11194823.G.A | rs11672123 | 0 | G | A | 0.05 | 0 | 0.3 | 9.2E-01 |

**Fig S11.** Visualization of the 3 LD bins in region 624

(A). Heatmap of the correlation matrix in region 624 for the 8 variants kept after pruning, with SNPs ordered by LD bin (A.). Panel

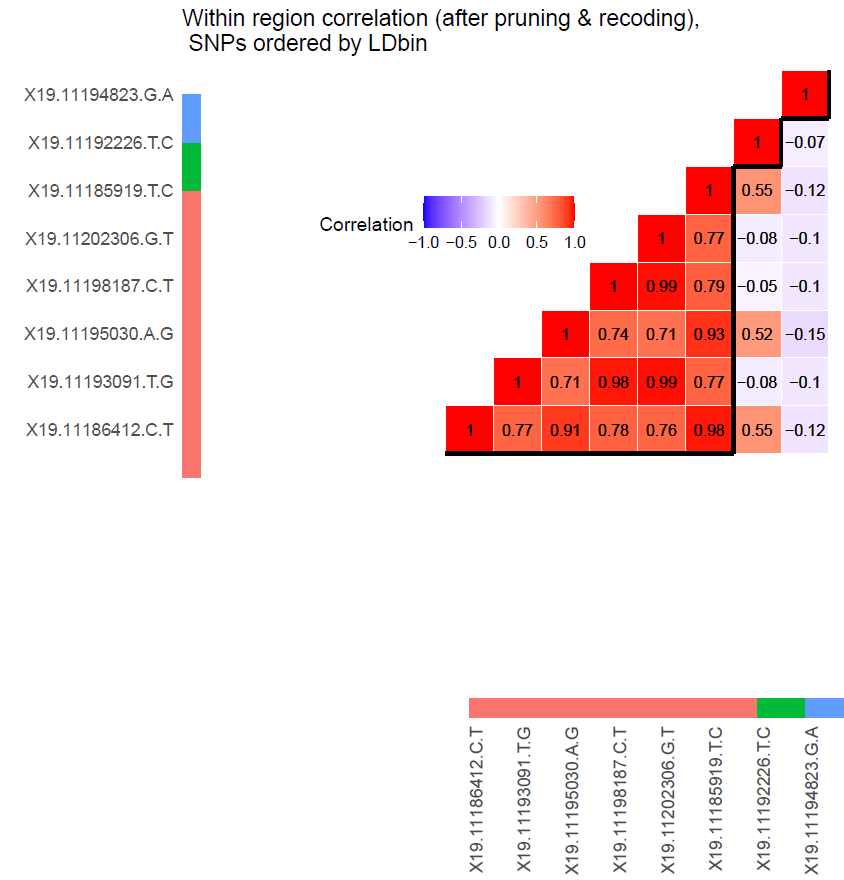

(B.) Position of the 8 SNPs (X axis) according to the bins they are assigned to (Y axis); the other SNPs shown in grey are the SNPs part of the same bins but pruned out because of high LD with other kept for the analysis.

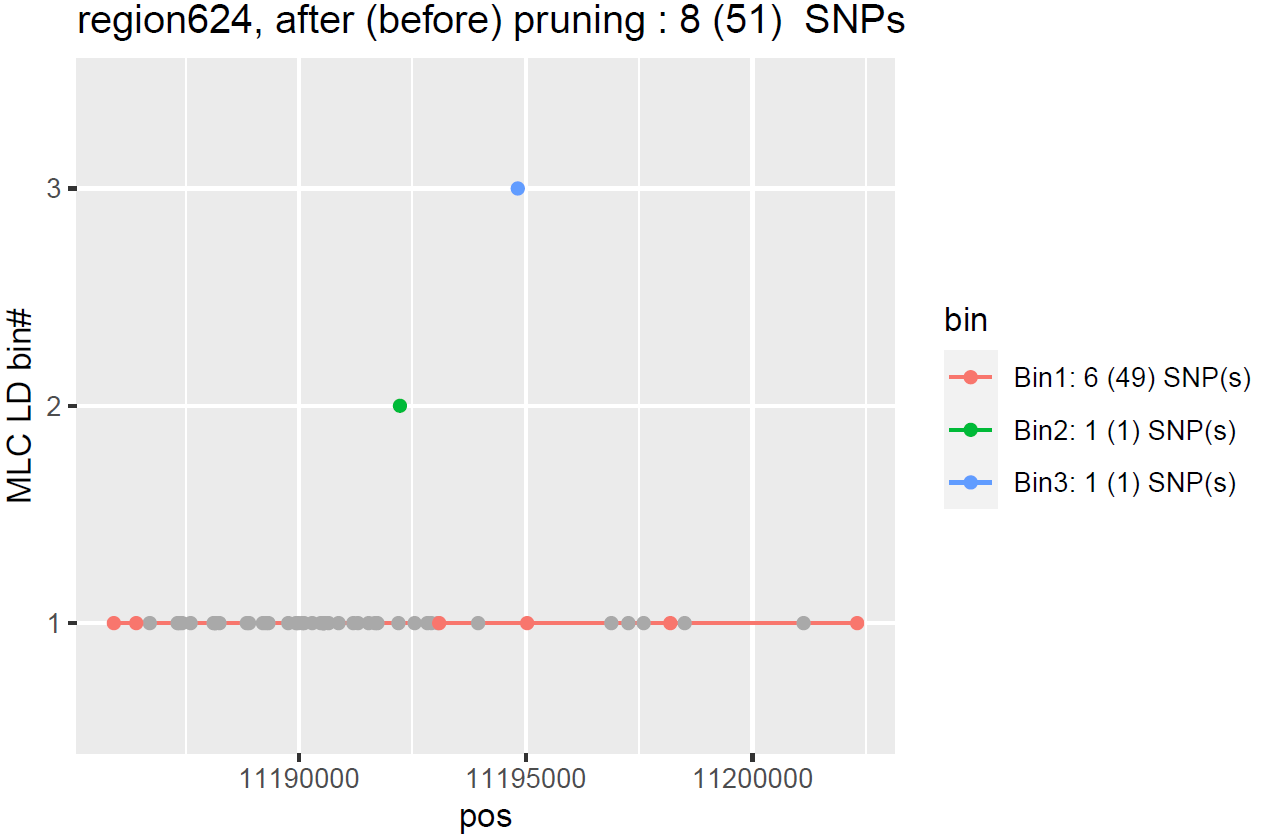

#### 3.4 List of the participants of the DCCT/EDIC Research Group (as of January 1, 2023)

*Study Chairpersons* – D.M. Nathan (chair), R. Gubitosi-Klug (vice-chair); *Past*: O. Crofford, B. Zinman; *Deceased*: S. Genuth

*Editor, EDIC Publications* – D.M. Nathan

**Clinical Centers**

Case Western Reserve University – *Current*: R. Gubitosi-Klug, L. Mayer, J. Wood, G. Greanoff, D. Miller, M. Novak, S. Pendegast, S. Rath, L. Singerman, D. Weiss, H. Zegarra; *Past*: E. Brown, P. Crawford, M. Palmert, P. Pugsley, J. Quin, S. Smith-Brewer; *Deceased*: W. Dahms, S. Genuth, J. McConnell

Weill Cornell Medical College – *Current*: N.S. Gregory, R. Hanna, R. Chan, S. Kiss, A. Orlin, M. Rubin; *Past*: S. Barron, B. Bosco, D. Brillon, S. Chang, A. Dwoskin, M. Heinemann, L. Jovanovic, M.E. Lackaye, T. Lee, B. Levy, V. Reppucci, M. Richardson; *Deceased*: R. Campbell

Henry Ford Health System – *Current*: A. Bhan, J.K. Jones, D. Kruger, P.A. Edwards, S. Mukhashen; *Past*: E. Angus, A. Galprin, M. McLellan, H. Remtema, A. Thomas; *Deceased*: J.D. Carey, F. Whitehouse

International Diabetes Center – *Current*: R. Bergenstal, S. Dunnigan, M. Johnson, A. Carlson, L. Thomas; *Past*: R. Birk, P. Callahan, G. Castle, R. Cuddihy, M. Franz, D. Freking, L. Gill, J. Gott, K. Gunyou, P. Hollander, D. Kendall, J. Laechelt, S. List, G. Matfin, W. Mestrezat, J. Nelson, B. Olson, N. Rude, M. Spencer; *Deceased*: D. Etzwiler, K. Morgan

Joslin Diabetes Center – *Current*: L.P. Aiello, E. Golden, P. Arrigg, R. Beaser, J. Cavallerano, R. Cavicchi, O. Ganda, O. Hamdy, T. Murtha, D. Schlossman, S. Shah, G. Sharuk, P. Silva, P. Silver, M. Stockman, J. Sun, E. Weimann; *Past*: V. Asuquo, L. Bestourous, A. Jacobson, R. Kirby, L. Rand, J. Rosenzwieg, H. Wolpert

Massachusetts General Hospital – *Current*: D.M. Nathan, M.E. Larkin, R. Azevedo, R. Bartholomew, K. Chu, J. Heier, A. Joseph, A. Leong, C. Shah, N. Thangthaeng; *Past*: E. Anderson, H. Bode, S. Brink, M. Cayford, M. Christofi, C. Cornish, D. Cros, S. Crowell, L. Delahanty, A. deManbey, K. Folino, S. Fritz, C. Gauthier-Kelly, J. Godine, L. Gurry, C. Haggan, K. Hansen, F. Leandre, P. Lou, J. Lynch, K. Martin, C. McKitrick, D. Moore, D. Norman, M. Ong, E. Ryan, C. Stevens, C. Taylor, D. Zimbler

Mayo Clinic – *Current*: A. Vella, A. Zipse, A. Barkmeier; *Past*: B. French, M. Haymond, J. Mortenson, J. Pach, R. Rizza, L. Schmidt, W.F. Schwenk, R. Woodwick, G. Ziegler; *Deceased*: R. Colligan, A. Lucas, F.J. Service, B. Zimmerman

Medical University of South Carolina – *Current*: H. Karanchi, L. Spillers, J. Fernandes, K. Hermayer, K. Lee, T. Lyons, M. Nutaitis; *Past*: A. Blevins, M. Bracey, S. Caulder, J. Colwell, S. Elsing, A. Farr, S. Kwon, D. Lee, P. Lindsey, M. Lopes-Virella, L. Luttrell, R. Mayfield, J. Parker, N. Patel, C. Pittman, J. Selby, J. Soule, M. Szpiech, T. Thompson, D. Wood, S. Yacoub-Wasef

Northwestern University – *Current*: A. Wallia, M. Hartmuller, M. El Muayed, M. Gill, A. Lyon, R. Mirza; *Past*: D. Adelman, S. Colson, M. Molitch, B. Schaefer

University of California, San Diego – *Current*: S. Mudaliar, G. Lorenzi, O. Kolterman, M. Goldbaum; *Past*: T. Clark, M. Giotta, I. Grant, K. Jones, R. Lyon, M. Prince, R. Reed, M. Swenson; *Deceased*: G. Friedenberg

University of Iowa – *Current*: W.I. Sivitz, B. Vittetoe; *Past*: M. Bayless, C. Fountain, R. Hoffman, J. Kramer, J. MacIndoe, N. Olson, H. Schrott, L. Snetselaar, T. Weingeist, R. Zeitler

University of Maryland – *Current*: R. Miller, S. Johnsonbaugh; *Past*: M. Carney, D. Counts, T. Donner, J. Gordon, M. Hebdon, R. Hemady, B. Jones, A. Kowarski, R. Liss, S. Mendley, D. Ostrowski, M. Patronas, P. Salemi, S. Steidl

University of Michigan – *Current*: W.H. Herman, R. Pop-Busui, C.L. Martin, P. Lee,  J. W. Albers, E.L. Feldman; *Past*: N. Burkhart, D.A. Greene, T. Sandford, M.J. Stevens; *Deceased*: J. Floyd

University of Minnesota – *Current*: A. Bantle, J. Bantle, M. Rhodes, D. Koozekanani, S. Montezuma, J. Terry; *Past*: N. Flaherty, F. Goetz, C. Kwong, L. McKenzie, M. Mech, J. Olson, B. Rogness, T. Strand, J. Terry, R. Warhol, N. Wimmergren

University of Missouri – *Current*: D. Hainsworth, S. Hitt, A. Jarvis; *Past:* D. Goldstein; *Deceased*: J. Giangiacomo

University of New Mexico – *Current*: D.S. Schade, A. Bancroft, R.B. Avery, M.R. Burge, J.E. Chapin, A. Das, L.H. Ketai; *Past*: J.L. Canady, D. Hornbeck, C. Johannes, J. Rich, M.L Schluter

University of Pennsylvania – *Current*: M. Schutta, P.A. Bourne, A. Brucker; *Past*: S. Braunstein, B.J. Maschak-Carey, S. Schwartz; *Deceased*: L. Baker

University of Pittsburgh – *Current*: T. Costacou, F. Toledo, T. Orchard, B.A. Coonrod; *Past*: D. Becker, L. Cimino, B. Doft, D. Finegold, K. Kelly, L. Lobes, D. Rubinstein, N. Silvers, T. Songer, D. Steinberg, L. Steranchak, J.Wesche; *Deceased*: A. Drash

University of South Florida – *Current*: J.I. Malone, A. Morrison, H. Rodriguez, J. O’Brian, P.R. Pavan; *Past*: L. Babbione, M.L. Bernal,T.J. DeClue, N. Grove, D. McMillan, H. Solc, E.A. Tanaka, J. Vaccaro-Kish

University of Tennessee – *Current*: S. Dagogo-Jack, R. Wilson, S. Huddleston; *Past*: M. Bryer-Ash, E. Chaum, A. Iannacone, H. Lambeth, D. Meyer, S. Moser, M.B. Murphy, A. Patel, H. Ricks, S. Schussler, C. Wigley, S. Yoser; *Deceased*: A. Kitabchi

University of Texas Southwestern Medical Center – *Current*: P. Raskin, L. Jordan, B. Shao, YG. He, E. Mendelson, RL. Ufret-Vincenty; *Past*: M. Basco; *Deceased*: S. Cercone, S. Strowig

University of Toronto – *Current*: B.A. Perkins, A. Barnie, N. Bakshi, M. Brent, R. Devenyi, K. Koushan, M. Mandelcorn, D. Olegario, F. Perdikaris; *Past*: D. Daneman, R. Ehrlich, S. Ferguson, A. Gordon, L. Leiter, K. Perlman, S. Rogers, L. Tuason, B. Zinman

University of Washington – *Current*: I. Hirsch, X. Averkiou, I.H. de Boer, L. Olmos de Koo; *Past*: S. Catton, R. Fahlstrom J. Ginsberg, J. Kinyoun, J. Palmer, L. Van Ottingham

University of Western Ontario – *Current*: C. McDonald, M. Driscoll, J. Bylsma, T. Sheidow; *Past*: W. Brown, C. Canny, P. Colby, S. Debrabandere, J. Dupre, J. Harth, I. Hramiak, M. Jenner, J. Mahon, D. Nicolle, N.W. Rodger, T. Smith

Vanderbilt University – *Current*: M. May, T. Marksbury, T. Adkins, A. Agarwal, C. Lovell; *Past*: S. Feman, J. Lipps Hagan, R. Lorenz, R. Ramker; *Deceased*: L. Survant

Washington University, St. Louis – *Current*: A. Brown, N.H. White, E. Hoffman; *Past*: L. Levandoski; *Deceased*: I. Boniuk, J. Santiago

Yale University – *Current*: W. Tamborlane, J. Sherr, P. Gatcomb, K. Stoessel; *Past*: J. Ahern

Albert Einstein – *Past*: J. Brown-Friday, J. Crandall, H. Engel, S. Engel, H. Martinez, M. Phillips, M. Reid, H. Shamoon, J. Sheindlin

**Clinical Coordinating Center**

Case Western Reserve University – *Current*: R. Gubitosi-Klug, L. Mayer, K. Farrell; *Past*: C. Beck, P. Gaston, M. Palmert, J. Quin, R. Trail; *Deceased*: W. Dahms, S. Genuth

**Data Coordinating Center**

George Washington University, The Biostatistics Center – *Current*: J. Lachin, I. Bebu, B. Braffett, J. Backlund, M. Bott, L. Diminick, L. El ghormli, X. Gao, S. Ho, D. Kenny, K. Klumpp, M. Lin, V. Trapani; *Past*: K. Anderson, K. Chan, P. Cleary, A. Determan, L. Dews, W. Hsu, P. McGee, H. Pan, B. Petty, D. Rosenberg, B. Rutledge, W. Sun, S. Villavicencio, N. Younes; *Deceased*: C. Williams

**National Institute of Diabetes and Digestive and Kidney Disease**

National Institute of Diabetes and Digestive and Kidney Disease Program Office – *Current*: E. Leschek; *Past*: C. Cowie, C. Siebert

**EDIC Core Central Units**

Central Biochemistry Laboratory (University of Minnesota) – *Current*: M. Steffes, A. Karger, J. Seegmiller, V. Arends; *Past*: J. Bucksa, B. Chavers, A. Killeen, M. Nowicki, A. Saenger

Central ECG Reading Unit (Wake Forest School of Medicine) – *Current*: E.Z. Soliman, M. Barr, C. Campbell, S. Hensley, J. Hu, L. Keasler, Y. Li, T. Taylor, Z.M. Zhang; *Past*: Y. Pokharel, R. Prineas

Central Ophthalmologic Reading Unit (University of Wisconsin) – *Current*: B. Blodi, R. Danis, D. Lawrence, H. Wabers; *Past*: M. Burger, M. Davis, J. Dingledine, V. Gama, S. Gangaputra, L. Hubbard, S. Neill, R. Sussman

Central Neuropsychological Reading Unit (NYU Long Island School of Medicine, University of Pittsburgh) – *Current*: A. Jacobson, C. Ryan, D. Saporito; *Past*: B. Burzuk, E. Cupelli, M. Geckle, D. Sandstrom, F. Thoma, T. Williams, T. Woodfill

### Supplementary Information 4. Evaluation of the effects of sample size and region size on computational time in an artificial dataset

In this section, we report estimates of computational runtime (elapsed time and CPU time) of *regscan* increase across a range of study sample sizes and region sizes (number of SNPs analyzed).

#### 4.1 Data generation

HAPGEN2^40^ was used to generate a balanced case-control design of 20,000 cases and 20,000 controls specified by a log-additive genetic model for the joint effects of five of the most frequent melanoma-associated missense variants in *MC1R* (**Table S19**). We simulated haplotypes and genotypes in 16q arm (GRCh38: 46280682p-end, 43Mb) from phased haplotypes for 156,673 biallelic SNPs (MAF>0.01) obtained from high-coverage whole genome sequencing (WGS) of 503 unrelated individuals of European ancestry from 1000G.^41^

**Table S19.** Specified odds ratio (ORs) for 5 *MC1R* causal variants in the log-additive multi-SNP model used to simulate 40,000 individuals (20,000 melanoma cases and 20,000 controls)

| *MC1R* causal variants | Chr:pos (GRCh38) | MAF^1^(%) | Minor/Major alleles | OR^2^ |
| --- | --- | --- | --- | --- |
| rs1805005 | 16:89919436 | 11.2 | T/G | 1.15 |
| rs2228479 | 16:89919532 | 6.9 | A/G | 1.22 |
| rs1805007 | 16:89919709 | 7.2 | T/C | 1.78 |
| rs1805008 | 16:89919736 | 6.2 | T/C | 1.43 |
| rs885479 | 16:89919746 | 7.0 | A/G | 1.42 |

^1^MAF in 503 individuals of European ancestry from whole genome sequenced individuals (30x) from 1000G

^2^Specified ORs for each of the 5 causal *MC1R* variants with MAF>0.01 in 1000G/EUR WGS individuals, as reported for melanoma risk^42^, using the minor allele as risk allele.

#### 4.2 Assessment of the computational efficiency

As described for the DCCT/EDIC analysis, we applied LD-based partitioning using BigLD/gpart^31,32^ to the 107,292 16q SNPs with MAF$\geq0.05$in 20,000 controls. We applied RegionScan to all the 2,394 regions identified by BigLD in the total sample (computational time, elapsed time 355.08 and CPU time 7405.07 minutes). We then randomly choose five regions with varying number of SNPs for analysis (after pruning): ~50 SNPs (region 1), ~100 SNPs (region 2), ~200 SNPs (region 3), ~400 SNPs (region 4), ~600 SNPs (region 5), and ran RegionScan again in N=20,000, N=10,000, N=5,000, N=2,500 randomly sub-sampled individuals. For each region, we report the elapsed time and CPU time (both in minutes) for each run of *regscan* in the total sample, and in the subsamples **(Fig S13**). Computational time assessments were all performed on a high-performance computer using a single node (each node has 40 CPUs and 202 GB RAM).

**Fig S12**. Distribution of region sizes (no. of SNPs analyzed) among the 2394 regions analyzed in the total sample size.

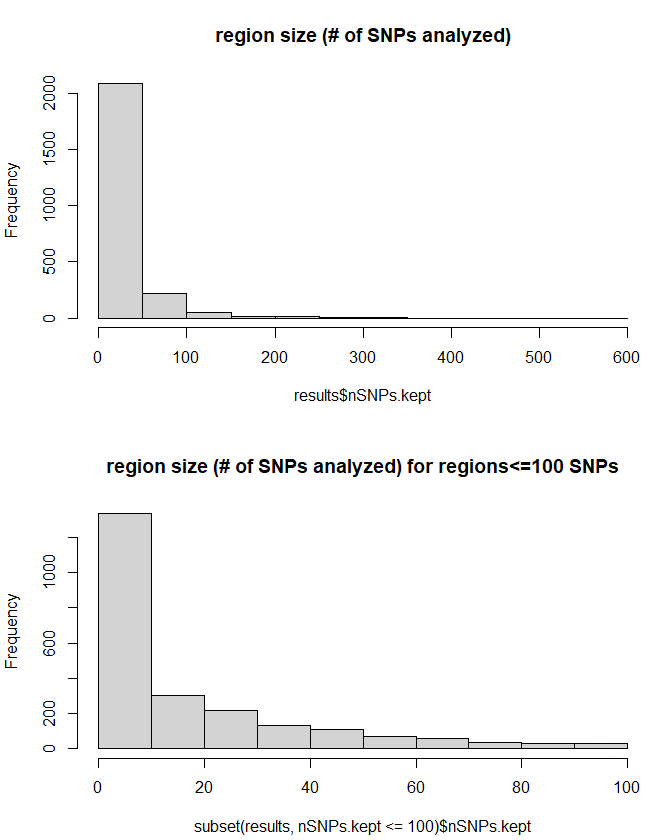

**Table S20.** Regions selected for the computational time assessment of *regscan* (ordered by number of SNPs kept after pruning on LD and removal of the aliases)

| Region # | Chr:start-end | Number of SNPs  after (before) pruning | Number of bins |
| --- | --- | --- | --- |
| Region 1 | 16: 50397624 - 50430199 | 50 (124) | 8 |
| Region 2 | 16: 80263964 - 80302523 | 100 (146) | 10 |
| Region 3 | 16: 80924545 - 81096217 | 201 (632) | 23 |
| Region 4 | 16: 82714946 - 82805368 | 305 (489) | 56 |
| Region 5^1^ | 16: 89605747 - 89926798 | 594 (1121) | 57 |

^1^Region5 includes all 5 causal variants specified under the melanoma-generating model.

**Fig S13.** Evaluation of *regscan* computational efficiency (*Y* axis: elapsed time, measured in minutes, log transformed) by region size (# of SNPs analyzed on the *X* axis) and sample size for the simulated melanoma binary outcome.

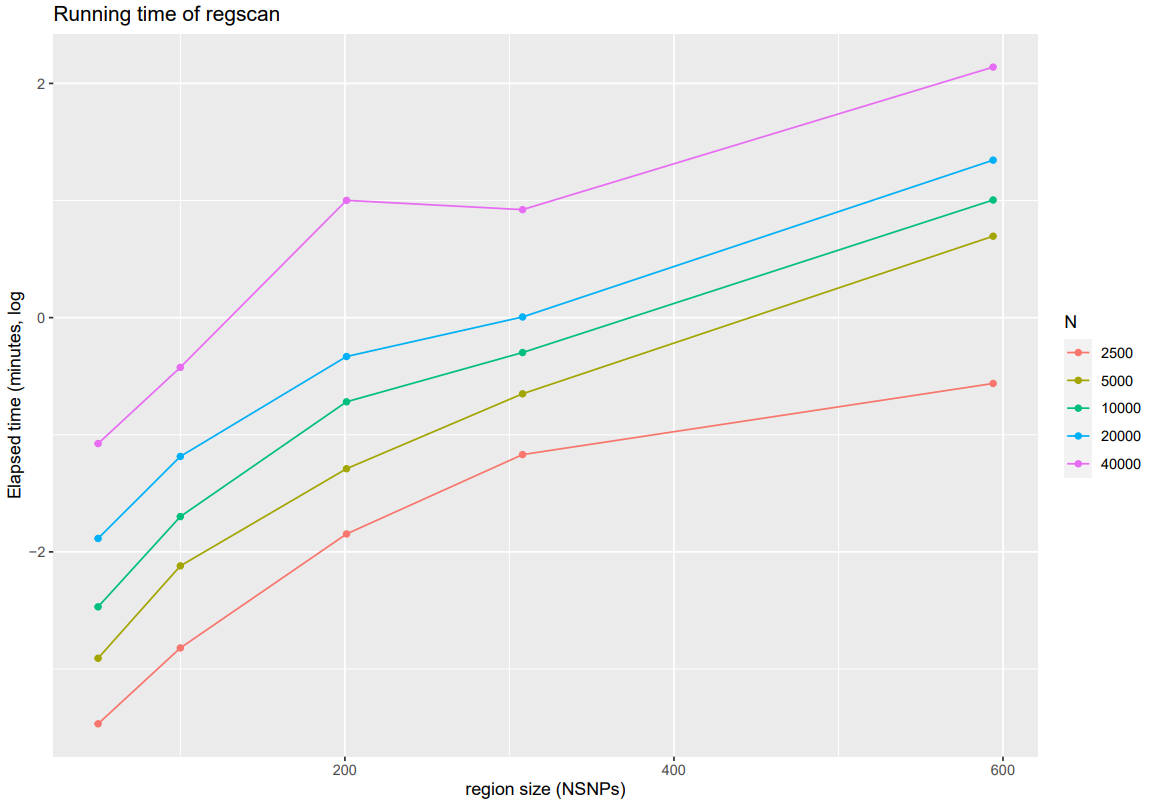

**Fig S14.** Evaluation of *regscan* computational efficiency (*Y* axis: elapsed time, measured in minutes) by region size (# of SNPs analyzed on the *X* axis) and sample size for the simulated melanoma binary outcome.

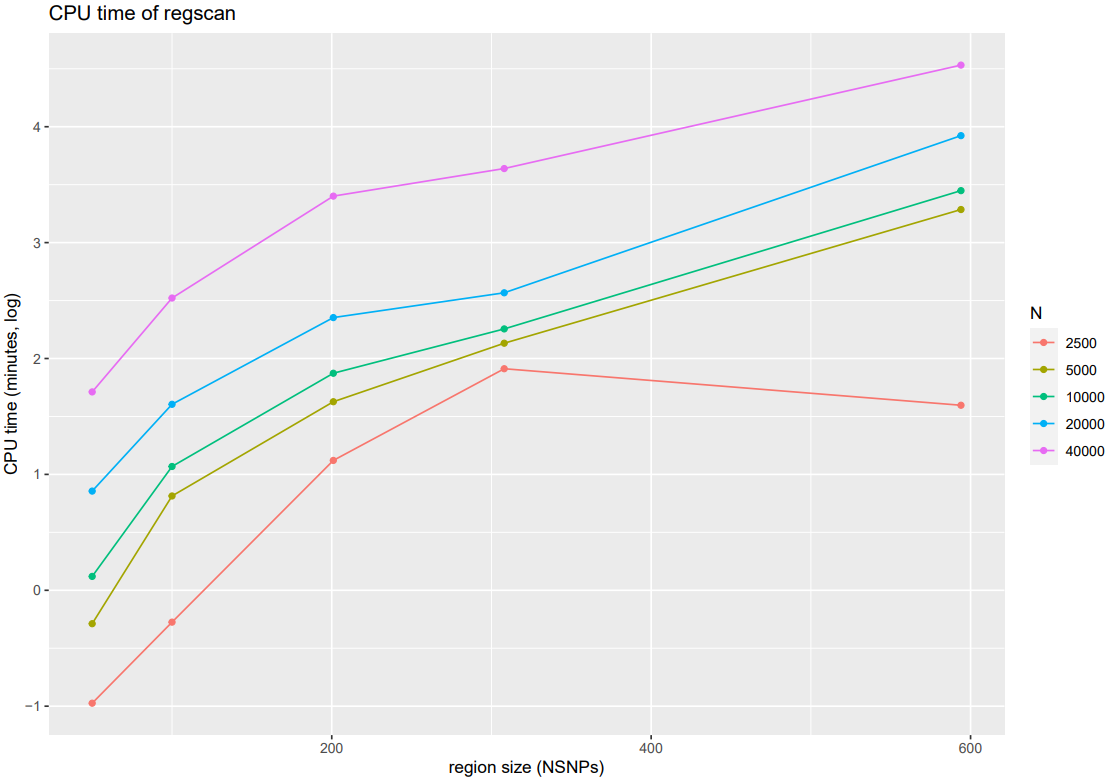
